# Rethinking the movement ecology of Andean bears: temperature-driven cathemerality and seasonal space-use cycles

**DOI:** 10.64898/2026.05.11.720697

**Authors:** Francisco X. Castellanos, David Jackson, Stefano Mezzini, Jorge Brito, Armando Castellanos

## Abstract

**Background:** The Andean bear (*Tremarctos ornatus*), South America’s only ursid, is one of the world’s most elusive large mammals, making movement data collection exceptionally rare. Addressing this gap, we present the largest telemetry dataset ever assembled, spanning 19 individuals tracked across three Ecuadorian National Parks over two decades, paired with a novel analytical approach.

**Methods:** We integrated Continuous-Time Movement Models (CTMM), Auto-correlated Kernel Density Estimates (AKDEs), Hidden Markov Models (HMM) and a diel niche theoretical framework to mitigate biases previously unaccounted for the species in telemetry studies. Fine-scale AKDEs and non-linear movement metrics were calculated to understand seasonal space use and movement behaviors. Speed and diffusion from CTMM and behavioral states from HMM were modelled with environmental covariates to investigate which conditions shape diel and seasonal activity.

**Results:** Population mean home range was 138.2 km^2^ (95% Confidence Intervals 78.7–225.5), with males (239.8 km^2^, 182.8–307.5), significantly exceeding females (58.5 km^2^, 35.5–90.3). Notably, three females exhibited ranges comparable to some males. Weekly and monthly AKDEs uncovered cyclic home range dynamics potentially driven by resource availability, with contractions around corn harvests and berry seasons, and expansions during páramo transitions. Decoupling speed from diffusion rates showed region-specific behaviors: intensive patch exploitation in Llanganates, broad exploratory ranging in Cayambe–Coca, and suppressed female locomotion in Cotacachi–Cayapas. Statistical analyses identified temperature as a key diel modulator and precipitation as the seasonal driver. Foraging probability increased between 2:00–6:00, large displacements between 7:00–14:00, and nocturnal movement rose significantly under colder conditions. Across diel hypothesis frameworks, bears were classified as cathemeral rather than strictly diurnal, corroborated by camera-trap records from Colombia, Ecuador, and Peru.

**Conclusions:** We propose a cathemeral diel phenotype that responds to thermal fluctuations and situates Andean bears within a broader ursid context of thermoregulatory niche plasticity. This dataset reveals unprecedented resolution of regional and sex specific behaviors that will facilitate and accelerate comparative studies in rapidly changing Andean landscapes. By releasing this long-term dataset as an open resource, we provide a foundation for climate-resilient conservation strategies. More broadly, we advocate for data democratization and invite collaboration.

## 1 Background

The Andean bear *Tremarctos ornatus* is a large elusive mammal, and the only extant bear species native to South America [1]. Its distinctive facial markings, often forming spectacle-like shapes, vary from individual to individual and can change through their lifespan [1–3], inspiring the common name “spectacled bear”. This species ranges from western Venezuela to Southern Bolivia and a controversial distributional boundary in Argentina [1, 4]. The species occupies a diverse range of habitats including páramo and Puna high grasslands, subtropical and tropical montane forests, dry forests, and even shrubby coastal deserts [1, 3].

In Ecuador, Andean bear populations are concentrated along the Andean mountain belt, between elevations of 700 and 4,900 m.a.s.l. [1]. Subpopulations exist in both the western and eastern foothills, spanning the Chocó bioregion and the upper Amazon basin, respectively [5]. Nevertheless, geographical barriers seem to represent no obstacle to gene flow, with studies revealing the existence of a single genetically homogenous population [6, 7].

Although research on Andean bears lags behind studies of other Ursids, recent work has contributed important insights mostly through noninvasive and incidental sampling, such as: population genetics [6, 8–11], occupancy models via camera trap data [12, 13], maternal and kinship relations [14–17], human-bear conflict [18], and parasitology [19, 20]. Despite the low capture success rate, collar tagging of over 20 specimens has been achieved in Ecuador [21, 22], two in Colombia [23, 24], two in Bolivia [25] and one in Peru [26]. From the collected data, analyses have largely been limited to home range estimation [22–24, 27] and used as proxies to understand their behavior. These studies relied on traditional estimators such as Minimum Convex Polygon (MCP), Kernel Density Estimate (KDE), and Local Convex Hull (LoCoH), without accounting for bias sources such as irregular sampling regimes, small effective sample sizes (ESS; i.e., number of home range crossings), and unmodeled autocorrelation in Global Position System (GPS) datasets [28–30]. Moreover, these methods assume each fixed position is Independent and Identically Distributed (IID), leading to negatively biased estimates when this assumption is violated. Mitigating these biases is particularly relevant as the availability of GPS data increases.

GPS collars enable fine-scale movement and behavioral data collection across temporal resolutions, but their application in Andean bear studies remains limited, largely due to the logistical and economic challenges of capture and collar-tagging, and the technological intricacies caused by GPS tracking in densely forested environments. The largest home range study, published more than a decade ago, relied on VHF data [22, 27], and little progress has been made in understanding the movement ecology of the species since then. Only two GPS-based studies in Colombia have been published in the last five years; together, they describe home ranges and core areas for two bears, with straight-line displacements estimated for only one individual [23, 24].

Diel activity patterns remain poorly understood. Camera trap and telemetry data from Chingaza Natural National Park in Colombia [31], and in the Apolobamba range in Bolivia suggest primarily diurnal movement [25]. However, Peyton B. noted day and night activity in cloud forests through extensive field research [33]. Similarly, anecdotal reports from Cotacachi-Cayapas National Park, and telemetry studies in the Maquipucuna Biological Reserve in Ecuador [34] note activity concentrated between 06:00–18:30 with short rest bouts in the afternoon, and nocturnal movements likely driven by low temperatures and high precipitation rates (Castellanos A. pers. comm.).

Critical knowledge gaps persist in fine-scale movement patterns and diel behaviors, which are essential for understanding diel niche partitioning, environmental drivers of movement, and individual variation within and between populations. This scenario calls for updated, robust, and comprehensive studies utilizing modern telemetry, computational technologies, alongside modelling techniques designed to mitigate bias sources [28].

After 20 years of intensive field work, tracking, and data collection, we present the largest telemetry dataset of an Andean bear population across three regions of the Ecuadorian Andes. Using high resolution GPS data from nine Andean bears inhabiting high elevation grasslands, and reevaluating VHF data from ten individuals [22], we applied novel computational and statistical methods to estimate population-level home range and core area sizes, non-linear movement metrics, and infer environmentally-driven behavioral responses. The availability of these new tracking units enables an unbiased examination of diel and seasonal dynamics under a robust framework that incorporates environmental data to evaluate drivers of movement and behavioral change. This work provides a novel, data-rich framework for assessing movement ecology in this emblematic Neotropical species, highlights the importance of democratizing data, and contributes to informed conservation planning while advocating for open ecological data to support collective research efforts. By releasing this dataset, we enable cross-country comparisons and collaborative conservation planning, demonstrating how democratizing long-term ecological data can accelerate both scientific progress and conservation outcomes.

## 2 Methods

### 2.1 Study areas

#### 2.1.1 Cotacachi-Cayapas National Park

This northernmost National Park (NP) covers 2,436.38 km^2^ across the provinces of Esmeraldas and Imbabura [35]. Elevations range from 30 to 4,939 m.a.s.l. [35]. Its western slopes fall within the Chocó ecoregion—one of the planet’s ten most important biological hotspots [36]. Three vegetation zones are present from lowest to highest: cloud forest, high montane forest and páramo. Above 2,000 m.a.s.l., *Cecropia* is common, along with Lauraceae species such as *Nectandra* and *Ocotea* [37]. In cloud forests, bears obtain protein and fiber from *Chusquea*, a bamboo-like species which makes up 75% of their daily diet [37]. In the páramo, Andean blueberries locally known as mortiño (*Vaccinium floribundum*) are an essential seasonal fruit in the diet of the Andean bear. Fructification occurs in two peaks: the highest from September to December and a secondary from January to April (Efrain Freire pers. comm.). The park experiences two climate types: Equatorial high mountain and Equatorial mesothermic semi-humid, with average annual temperatures fluctuating between 12°C and 18°C [35, 38]. Rainfall is concentrated between October and May, followed by a dry season from June to September [38]. Human activities, particularly maize production in Intag, drive conflict between bears and farmers [37]. Burning, overgrazing, and illegal mining have also fragmented and reduced *Polylepis* forests [35].

#### 2.1.2 Cayambe-Coca National Park

It encompasses an area of 4,083 km^2^, with altitudinal gradients from 600 to 5,790 m.a.s.l., and spans the provinces of Pichincha, Imbabura, Napo and Sucumbíos [39]. The NP includes both the Andean and Amazon regions, with vegetation ranging from high montane forest to páramo [40], and two zoogeographic floors: temperate and high Andean [41]. It has a delimited rainy season from June to August, dry from September to January, and intermittent rains during the rest of the year [38, 42]. The climate is classified as Equatorial mesothermal and semihumid [38, 42]. Mean temperatures vary with elevation but average around 8°C, with maximum temperatures rarely exceeding 20°C and minimums often approaching 0°C [39]. The park’s highlands form a vital hydrological reservoir, including several wetland systems designated as the Ñucanchi Turupamba Ramsar site [39], one of the 19 Ramsar sites present in Ecuador.

These wetlands are critical for biodiversity, water provision and climate change mitigation [39]. The NP hosts over 100 endemic plant species, and páramo remnants of *Polylepis* forests serve as refuge for Andean wildlife [39]. Several species of bromeliads like “achupallas” (*Puya* spp., *Greigia* spp.) constitute a key component of Andean bear diet, as they heavily feed on their meristematic tissues and leaf bases. These plants are scattered throughout the páramo in patches and are abundant year-round [43]. Additionally, berries of the genus *Rubus*, *Escallonia myrtilloides*, and *Pernettya prostrata* are available during the rainy season [43]. However, because extensive cattle ranching overlaps with the natural habitat of bears, conflict with humans is common within the Kichwa indigenous community of Oyacachi, primarily due to cattle predation [44].

#### 2.1.3 Llanganates National Park

The southernmost study site covers 2,197 km^2^, extending from the Andean to the Amazon region in the provinces of Cotopaxi, Tungurahua, Napo and Pastaza [45]. Within the park is the Llanganati Complex considered a Ramsar site [45]. The NP is found in the province of Tungurahua, within the temperate zoogeographic floor [41], where altitudes range from 1,200 to 4,570 m.a.s.l. [45]. The prevailing climates are Equatorial high montane and mesothermic semi humid with annual average temperatures that fluctuate between 12°C and 22°C. Vegetation zones include cloud forest, high montane forest and páramo [40]. Rainfall is strongly seasonal, with a rainy period from June to August, a dry season from September to January and intermittent rains during the rest of the year [46]. Over 800 vascular plant species have been identified throughout the park [45]. In the páramo, berries and achupallas are available with seasonal resemblance to the Cotacachi-Cayapas NP (Efrain Freire pers. comm.). Agricultural and livestock activities in cloud forest areas have caused habitat fragmentation. Maize is the primary crop harvested from January to May, and frequent bear incursions into these fields have led to conflict with local farmers (Edgar Martínez pers. comm.).

### 2.2 Capture and GPS collar tagging

Over a 9-year period, we captured and tagged 3 males and 6 females in the Cayambe-Coca and Llanganates NPs, representing different life stages (subadult, adult, and long-lived; Figure 1, Table 1).

**Fig. 1.**
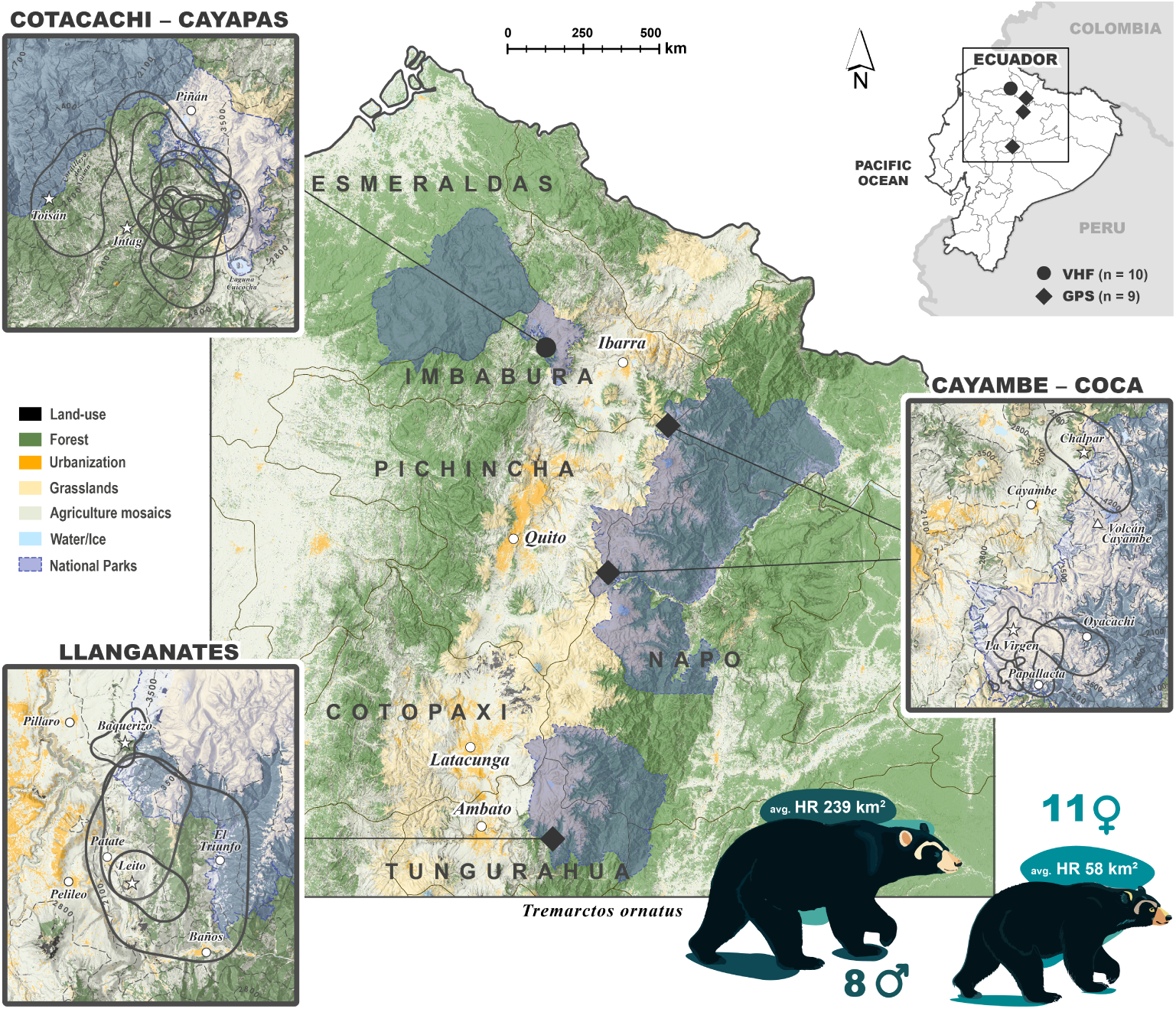
Telemetry study sites of Andean bears in Ecuador. Topographic map overlaid with land cover data from MapBiomas Ecuador (2023 [47]). Study sites where VHF and GPS data was generated are shown with a circle or diamonds, respectively. Zoomed-in maps show 95% AKDEs of bears tracked in each region with a grey continuous line. Elevation contours are shown with dashed lines and were generated in ArcGis Pro with Digital Elevation Model data downloaded with the elevatr R package v0.99.0 [48]. Major cities and important localities to this work are shown with a white circle whereas the stars show approximate capture localities. Sex-based population average home range (HR) is shown with each bear

**Table 1.** Animal fix data. Andean bears tracked with VHF and GPS collars in three National Parks: Cayambe-Coca (CCNP), Llanganates (LLNP), and Cotacachi-Cayapas (CCYNP). Individuals from CCYNP were marked with VHF collars [22].

| Name | Sex | Life stage | Fixes | Tracking period | # months | Region |
| --- | --- | --- | --- | --- | --- | --- |
| Chalpar | male | subadult | 7,622 | Aug 2019 – May 2022 | 34.2 | CCNP |
| Daniela | female | adult | 3,960 | Oct 2019 – Sep 2020 | 11.7 | CCNP |
| Matias | male | adult | 3,219 | Apr 2018 – Dec 2020 | 33.1 | CCNP |
| Paya | female | adult | 819 | Jun 2017 – Jan 2018 | 7.8 | CCNP |
| Rebecca | female | adult | 887 | Sep 2015 – Mar 2016 | 6.8 | CCNP |
| Alejandra | female | subadult | 5,047 | May 2021 – Sep 2022 | 16.5 | LLNP |
| Armando | male | subadult | 1,915 | Mar 2023 – Dec 2023 | 9.7 | LLNP |
| DeeAnn | female | long-lived | 527 | Mar 2022 – Aug 2022 | 5.5 | LLNP |
| Lorena | female | subadult | 2,395 | Dec 2022 – Sep 2023 | 8.7 | LLNP |
| Amanda | female | adult | 200 | May 2003 – Mar 2005 | 22.4 | CCYNP |
| Carolina | female | subadult | 178 | Dec 2007 – May 2009 | 16.6 | CCYNP |
| Dolores | female | adult | 330 | Oct 2002 – Jan 2005 | 27.7 | CCYNP |
| Ezequiel | male | adult | 101 | Mar 2003 – Oct 2003 | 6.3 | CCYNP |
| Fernando | male | adult | 42 | Jan 2008 – Dec 2009 | 23.0 | CCYNP |
| Jaime | male | adult | 136 | Mar 2004 – Dec 2005 | 21.6 | CCYNP |
| Gilo | male | adult | 130 | Jan 2008 – Dec 2009 | 23.5 | CCYNP |
| Marjorie | female | adult | 249 | Oct 2000 – Jul 2003 | 32.7 | CCYNP |
| Pancho | male | subadult | 133 | May 2003 – Jun 2004 | 12.7 | CCYNP |
| Porraca | female | subadult | 513 | Jan 2002 – Dec 2005 | 47.4 | CCYNP |

Capture methods included Iznachi box traps, snare traps, and pursuit with trained dogs [21, 22, 49, 50]. Once captured, individuals were sedated via dart injection using an airsoft rifle (Daninject, USA). The immobilization mixture consisted of xylazine hydrochloride (0.3–2 mg/kg; Equiet, ERMA Labs, Cundinamarca, Colombia) and ketamine (3–8 mg/kg; Ket-A-100, Grupo Grandes, Quito, Ecuador [22]). Following immobilization, the bears were positioned in lateral decubitus, measured, assessed for health status, sampled for hair and blood, and fitted with an Iridium/GPS Vertex Lite collar (Vectronic Aerospace GmbH, Berlin, Germany). GPS collars were programmed to record positions every two hours (12 fixes per day), until they detach naturally or were remotely released using a drop-off mechanism. When necessary, supplementary doses of ketamine hydrochloride (1–2 mg/kg intramuscular) were administered to maintain anesthesia until collar deployment was complete. Individuals were subsequently revived using the reversal agent yohimbine hydrochloride (0.1–0.25 mg/kg; Yohimbine Vet, Holliday-Scott S.A., Buenos Aires, Argentina [51]). Post procedure, proximity was maintained to confirm full recovery and good health.

### 2.3 Data processing and movement modeling

We analyzed movement data from nine GPS-tagged, and reanalyzed fifteen VHF-tagged individuals (Table 1, [22]). For VHF data, a fixed 81 m triangulation error was applied [22]. GPS location errors were quantified using calibration data and the root-mean-square user equivalent range error [52] in the R package ctmm v1.2.0 [53].

Based on delta corrected Akaike Information Criteria (ΔAICc), individual-specific error models were selected over population-level models (Additional file 1), revealing a horizontal error range of 1–32 m, and vertical of 2–348 m (Additional file 2; Table S1). Data was screened for unrealistic fixes by keeping those within the 2,000–5,000 m.a.s.l. altitudinal range and identifying minimum step-averaged speed outliers, typically marked as such if ≥0.4 m/s. These curated datasets, comprising 26,391 GPS and 2,012 VHF fixes, are available in the Movebank Data Repository [54]. Calibration data are not included because they were collected on private properties. However, R Data Serialization (RDS) objects containing the error models are available at https://github.com/Xacfran/andean bears.

Range residency was assessed by fitting semi-variance functions and inspecting variograms [55]. All GPS-collared bears exhibited range-resident behavior roughly after 3 months (Additional file 3, Fig. S1), whereas five VHF-tagged bears were excluded due to non-residency. Finally, we fit Continuous Time Movement Models (CTMM) by selecting the best-fit model for each individual via AICc. Models were fit to both the full datasets and the subsetted diurnal/nocturnal periods to account for diel variation. For details on error model selection, CTMMs and parameter estimates, see Additional file 1 and Additional file 2 (Table S2).

### 2.4 Home ranges and social interactions

We used the range distribution home range concept to underpin the possible future space use of the bears [56]. We hereafter use the terms “home range” as 95% of the utilization distribution (UD), “core area” as 50% of the UD, and “areas” to refer to both. To account for autocorrelation, small ESS, and irregular sampling, we used the optimally weighted area-corrected Autocorrelated Kernel Density Estimate (*wAKDE_c_* [57, 58]), hereafter “AKDE”, within ctmm. A key advantage of this framework was the ability to perform population-level inference; we estimated population mean AKDEs per region, sex, and life stage using a *χ*^2^ inverse Gaussian meta-analysis [59].

We visualized that large portions of UDs were shared by sympatric individuals. Given that direct observations of Andean bears have shed light on the interplay of complex social and hierarchical behaviors [17, 31], we examined potential social interplay by measuring home range overlap with the Bhattacharyya coefficient (BC) and estimated the probability of long-term encounters through the Conditional Distribution of Encounters (CDE [60]).

Hitherto, a temporal fine-scale understanding of bears’ home ranges has been unexplored in this species. To capture temporal variations, we estimated weekly AKDEs for GPS data using a lag bin width *dt* = 1 day (i.e., step size every 1 day), and monthly AKDEs for VHF data (*dt* = 7 days) with the window hr R script [61]. Importantly, weekly and monthly AKDE estimates reflect area shift intensity rather than home range values because AKDE estimations rely on stationarity assumptions; hence, large displacements within subsampled periods result in area overestimation.

For both datasets, we incorporated high-resolution satellite imagery using the basemaps v0.0.7 R package [62] to visually analyze, and describe individual behaviors across windows and regions. This detailed analysis is found in Additional file 1, and the animated movements at the Zenodo repository [63].

### 2.5 Behavioral states and diel niche analysis

We inferred behavioral states from GPS tracking data using Hidden Markov Models (HMM) to characterize movement persistence, activity budgets, and diel variation in relation to temperature and precipitation as independent variables. GPS points were annotated via the Environmental Data Automated Track Annotation System (Env-DATA [64]) with a bilinear interpolation, incorporating ERA5 2 m air temperature [65], and total precipitation [66]. Likewise, elevation was annotated from the Shuttle Radar Topography Mission at 1 arc-second, roughly 30 m (https://dwtkns.com/srtm30m/).

Because HMMs require regular sampling, we filtered GPS datasets into tracks representing at least 2 days of continuous locations with a maximum 9-hour gap and predicted missing positions with ctmm (11.8% of the final dataset; Additional file 2, Table S3). We used the R package momentuHMM v1.5.5 [67] to infer three a priori behavioral states: encamped, exploratory, and travelling (i.e., long displacements). None of the required initial model parameters exist for this species, thus, we iterated through 200 randomized parameter sets and selected the best-fit model with valid pseudo-residuals. Transition probabilities were modeled using weather covariates and cyclical time functions from the psych v2.4.6 package [68]. Diel cycles were best supported by temperature, while seasonal transitions were driven by precipitation. Detailed description of this process is provided in Additional file 1.

Mammalian diel research traditionally assigns species into single diel niches as fixed behaviors. In some ursids, however, evidence shows they display extreme diel plasticity based on sex, age [69], seasonality [70], and anthropogenic factors [69–71]. We investigated diel plasticity at a weekly-scale using the Diel.Niche v0.0.1.0 R package [72], which uses Bayesian model selection via Bayes factors to compare alternative categorical diel phenotypes. Weekly units were formed for each combination of individual, behavioral state, sex, and region, retaining *n* ≥ 20 GPS fixes per diel period category, yielding 426 weekly units across 7 individuals. Activity phenotypes were classified under three progressively more restrictive hypothesis frameworks: Traditional, General, and Selection to estimate phenotype probabilities across periods. See Additional file 1 for more details.

### 2.6 Speeds and diffusion rates

Traditional calculations of linear speeds and distances are scale-sensitive and biologically inaccurate as they tend to produce estimates biased by sampling frequency and measurement error [73]. Therefore, we calculated scale-insensitive continuous-time speeds and distances (CTSD [73]). Whole-tracking-period and diel speed estimates were restricted to GPS-tagged individuals, where sampling resolution was sufficient to resolve velocity autocorrelation, and yielded average speeds for seasonal inferences, and instantaneous speeds from diurnal and nocturnal subsets for a higher-resolution diel analysis.

#### 2.6.1 Fine-scale movement estimates

Because our sampling intervals exceeded velocity autocorrelation timescales, movement metrics at a daily temporal scale were rendered unreliable. Hence, we estimated monthly diffusion rates and average speeds for both GPS and VHF-tagged bears, as the coarser positional autocorrelation timescales governing diffusion are compatible with the sampling resolution of both tag types [74]. Diffusion, defined as the displacement variance over a finite period of time, integrates both movement rate and directional persistence, and therefore reflects broader-scale processes such as behavioral state switching, site fidelity, and exploratory phases [74].

We estimated weekly average speeds and diffusion rates using a sliding 4-week window and *dt* = 1 week by modifying the windows hr script [61]. CTMMs were fitted and selected for each window and diffusion rates were extracted from the model summary. Quality filtering retained only windows where diffusion degrees of freedom (DOF) ≥ 5, excluding infinite confidence intervals (CI), and removing IID models.

#### 2.6.2 Statistical tests

Population-level statistical comparisons across diel periods, sexes, ages, and regions were performed using meta-analyses in ctmm. Additionally, non-parametric tests to assess regional, sex, and diel differences in instantaneous speeds were performed via ggstatsplot v0.13.1 [75]: Mann-Whitney and Kruskal-Wallis. Effect sizes are reported as rank-biserial correlations (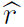).

Region-specific seasonal patterns in weekly speed and diffusion were modeled using Generalized Additive Models (GAM) in the mgcv R package [76] using REML smoothness selection. Each model included a sex and region interaction, a cyclic cubic spline for population-level and region-specific seasonality, and an individual-level random intercept.

Furthermore, to investigate the specific environmental conditions under which nocturnal movements were triggered, we tested the synergistic influence of temperature, precipitation, elevation, time of day, and seasonality on instantaneous speeds, by applying Hierarchical GAMs (HGAMs [77]). We used the bam function in mgcv, and specified a Gamma distribution with a log link, using the fREML method for efficient parameter estimation [78, 79]. The model incorporated cyclic cubic splines for diel and annual cycles and tensor product interactions to capture the joint covariate effects, and a correction for individual sampling interval bias. We accounted for individual variability using random intercepts and included sex as a fixed effect [77].

Model fit was assessed using deviance residuals. Residual-versus-linear-predictor plots showed structure consistent with the Gamma mean–variance relationship not fully capturing the spread of extreme instantaneous speeds, and the deviance residuals were right-skewed with a heavy upper tail (Additional file 3, Fig. S2). To ensure that this did not compromise our results, we evaluated the fitted conditional means directly through predictive checks, comparing predicted and observed speeds at the individual level. Even though the model under-predicted the fastest instantaneous speeds, it fully captured the overall distribution of speeds and the behavioral patterns present in the real data (Additional file 3, Fig. S2).

## 3 Results

### 3.1 Regional and sex-based area estimates

The population home range mean was 138.21 km^2^ (95% CI: 78.67–225.5). Males areas were significantly larger than those of females (*p <* 0.001). However, UD estimates revealed substantial individual variation. The females Porraca, Daniela and Alejandra had similar home ranges than Fernando and Ezequiel, and core areas similar to Jaime, Fernando, Matias and Ezequiel. For the remaining females, home ranges spanned from 20.4–90.3 km^2^, and core areas 3.3–18.8 km^2^, with the smallest values corresponding to Lorena and Rebecca, respectively (Fig. 2, Additional file 2, Table S4 and S5).

**Fig. 2.**
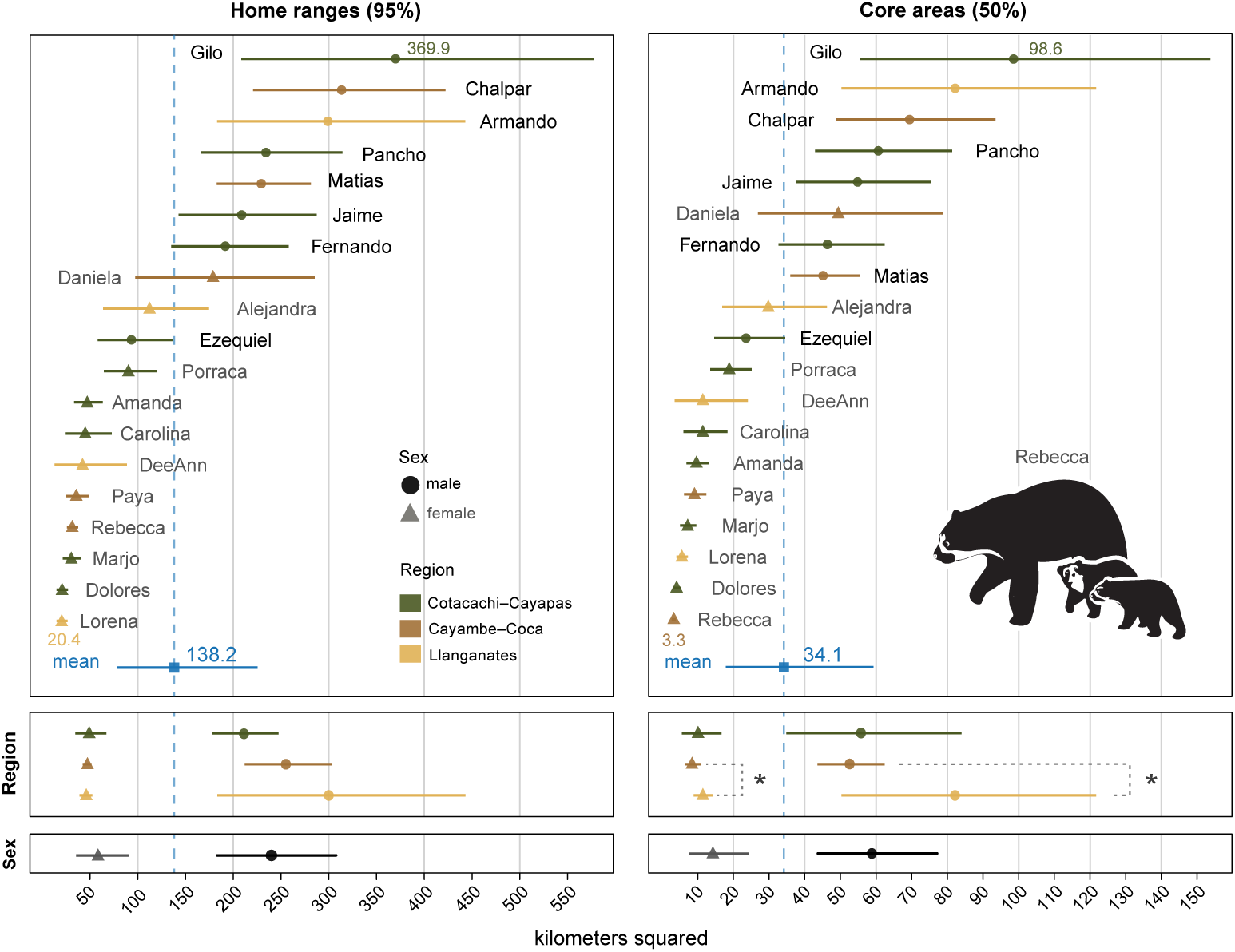
Home range and core area AKDE estimates of Andean bears in three Ecuadorian National Parks. Forest plots display individual and population-level averages and confidence intervals. Males are marked with a circle, females with a triangle and the population mean is shown with a blue dashed line. Population-level analyses and statistical significant testing were performed using a *χ*^2^ inverse Gaussian meta-analysis. The vector was generated by FXC from an illustration by Daniel Torres.

For males, the home range of Armando from Llanganates was the largest (299 km^2^), followed by Matias in Cayambe-Coca (255.3 km^2^), and an average of 211.12 km^2^ across four males in Cotacachi-Cayapas. In contrast, females exhibited the opposite trend (Fig. 2; Additional file 2, Table S6).

Regionally, the overall bear population showed no significant differences in home range sizes, nor did we find statistical differences between life stages (*p >* 0.05). Within sexes, significant differences were detected between the core areas of females from Cayambe-Coca (8.5 km^2^) and Llanganates (11.5 km^2^; p = 0.015), as well as between males from the same regions (52.64 km^2^ vs 82.19 km^2^) with p = 0.023 (Fig. 2; Additional file 2, table S6).

#### 3.1.1 Fine-scale home ranges

Seven– and thirty-day home range estimates revealed marked variation in movement strategies across individual Andean bears tied to sex, geography, and seasonal resource availability (Additional file 3, Fig. S3). During the dry season, between the months of January and May, bears within Llanganates occupied highly localized areas (Fig. 3), coinciding with the availability of corn (*Zea mays*) and corroborated by frequent visual records of bears near croplands (Additional file 3, Fig. S4). In contrast, bears in Cayambe-Coca displaced across expansive areas during the same period, suggesting divergent seasonal responses between landscapes (Fig. 3). Rainfall onset dramatically influenced space use. From late June to early October, Cayambe-Coca bears showed predominantly stationary behaviors with restricted ranges, punctuated by brief and intermittent excursions. In Llanganates, movement patterns reversed midyear; bears gradually expanded their home ranges and displayed fewer stationary periods through the second half of the year (Fig. 3).

**Fig. 3.**
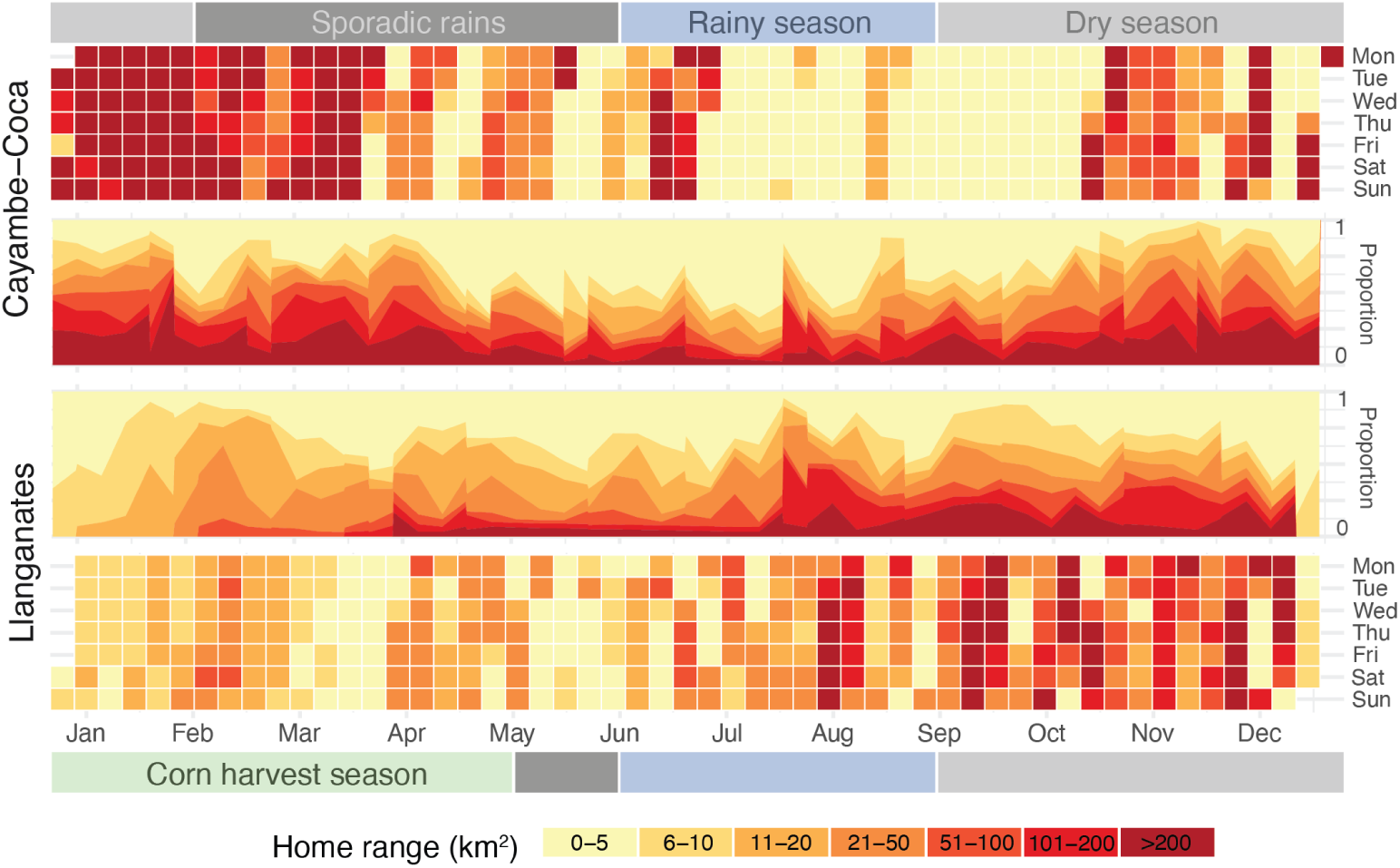
Daily, weekly, and monthly home range sizes. Heatmaps show aggregated counts of the most predominant home range size intervals in each day of the week per month color coded and described in the legend. Stacked areas accompany each heatmap displaying the relative contribution of each day of the week to the total events.

Male bears in Cayambe-Coca exhibited large and persistent displacements (*>*50 km^2^), while following defined routes that encompass multiple lakes and prominent geographic features (Additional file 1), similar to Armando in Llanganates. Notwithstanding, males in Cayambe-Coca exhibited more constrained movements than in other regions, and appeared to remain relatively stationary, from May to September, rather than travelling across their habitat (Additional file 1, Additional file 3, Fig. S3). Remarkably, Chalpar remained in a 3–21 km^2^ area at an altitude of 4,000 m.a.s.l. from September–October, close to the top of the inactive Cayambe volcano (Additional file 1).

Female movement patterns in all regions were more localized. For example, Rebecca and Paya exhibited high site fidelity (Additional file 1), reflecting a combination of habitat specialization and avoidance of human related risks near high traffic infrastructure. Similarly, in the Llanganates NP, DeeAnn and Lorena exhibited highly stable and compact home ranges (Additional file 1). Both individuals exhibited patterns of short distance displacements followed by extended stationary periods, likely driven by site-specific resource availability and habitat preferences.

Nevertheless, this was not the rule for the females. In Cayambe-Coca, multiple females demonstrated large-scale displacements with peaks (*>*50 km^2^) during March–June, and pronounced movement plasticity. Daniela, for example, expanded her range dramatically during seasonal pulses before contracting to microhabitats as small as 0.35 km^2^ for brief durations (Additional file 1; Additional file 3, Fig. S5).

In Cotacachi-Cayapas, large displacements were prevalent through April–June and mostly attributed to Carolina’s dispersal behavior (Additional file 3, Fig. S5). This oscillation between long distance movements and localized residency suggests a complex spatial strategy possibly linked to reproductive or foraging behavior. Together, these seasonal shifts reveal flexible movement strategies that respond to resource availability and landscape context. Detailed bear-by-bear movement profiles are presented in Additional file 1.

#### 3.1.2 Home range overlaps

Overlap estimates highlighted differences in shared space use between regions. In Llanganates, Armando and Alejandra exhibited the highest home range overlap (median BC = 0.78), followed by DeeAnn and Armando (0.64), and DeeAnn and Alejandra (0.54), indicative of shared resource zones. The CDE was located close to the center of Armando’s home range with a total area of 71.31 km^2^ (95% CIs: 42.28–107.79; Fig. 4). In Cayambe-Coca, overlaps were mostly lower than 0.15, suggesting strong spatial segregation. Matias and Daniela showed moderate overlap (0.38), consistent with their synchronous tracking periods. Matias and Paya overlapped 0.37 of their respective home ranges (Fig. 4; Additional file 1). The CDE area was 94.91 km^2^ (67.11–127.44), and centered on Matias’ home range, mostly encompassing the UDs of Paya, Rebecca, and the northwestern portion of Daniela’s (Fig. 4).

**Fig. 4.**
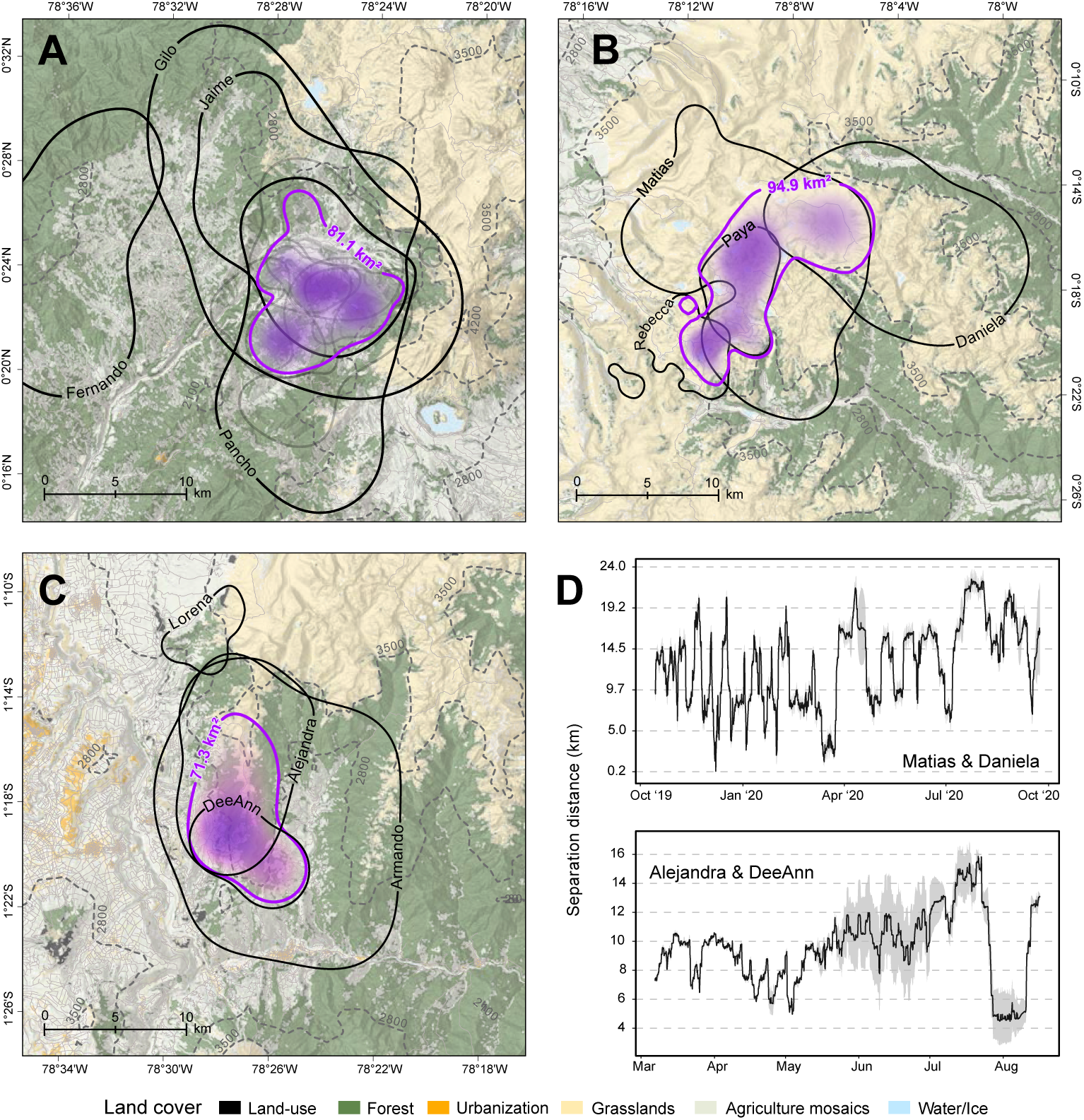
Home range overlap and Conditional Distribution of Encounters (CDE) across three Ecuadorian National Parks. Individual 95% AKDEs are overlaid on landcover and elevation data for: Cotacachi-Cayapas (A), Cayambe-Coca (B), and Llanganates (C). Purple polygons delineate the 95% CDE contours and the shaded raster represents the probability density function of the CDE. Roads and elevation contours are displayed with solid and dashed grey lines, respectively. (D) Pairwise separation distances in synchronously tracked individuals over time. Grey shaded areas indicate the 95% CIs.

A markedly different pattern emerged in Cotacachi-Cayapas, where all individuals UDs overlapped—except for Fernando’s—with a CDE measuring 81.13 km^2^ (65.11–98.88; Fig. 4). Notably, overlaps between and within sexes were substantial (*>*0.5; Additional file 2, Table S7), some as large as 0.82 in the case of Porraca and Pancho. Values lower than 0.5 are described in Additional file 1.

### 3.2 Quite diurnal, quite nocturnal

#### 3.2.1 Behavioral states

Behavioral state inference revealed the step lengths in the encamped state were 35.1 m (95% CI: 30.74–39.45), 180.02 m (161.74–198.3) during foraging, and 607.83 m (558.17–657.51) while travelling (Additional file 2, Table S8; Additional file 3, Fig. S5). Movement persistence was evident at the population-level in the travelling state, with males displaying more consistent directional movement at the individual level, except for Rebecca, whose movements were ballistics similar to males (Additional file 3, Fig. S5). On average, GPS–collared individuals spent approximately 32% of their sampling period encamped, 44% foraging, and 23% travelling long distances (Additional file 2, Table S8).

These states exhibited strong cyclical variation, capturing temperature as a critical diel behavioral modulator (Fig. 5A). Rising temperatures were positively associated with foraging and negatively with travel (Fig. 5B). Foraging stationary probabilities peaked at approximately 70% between 2:00 and 6:00. After 8:00, the probability of transitioning to travelling increased to roughly 60% and declined after 14:00. By 17:00, bears were more likely to transition into a prolonged encamped state, which waned after midnight as foraging resumed. In contrast, modeling the encamped state was statistically constrained; the extremely low transition probabilities from this state to others when the time covariate was fixed at noon (Additional file 3, Fig. S5), precluded a reliable estimation of stationary probabilities for this behavior.

**Fig. 5.**
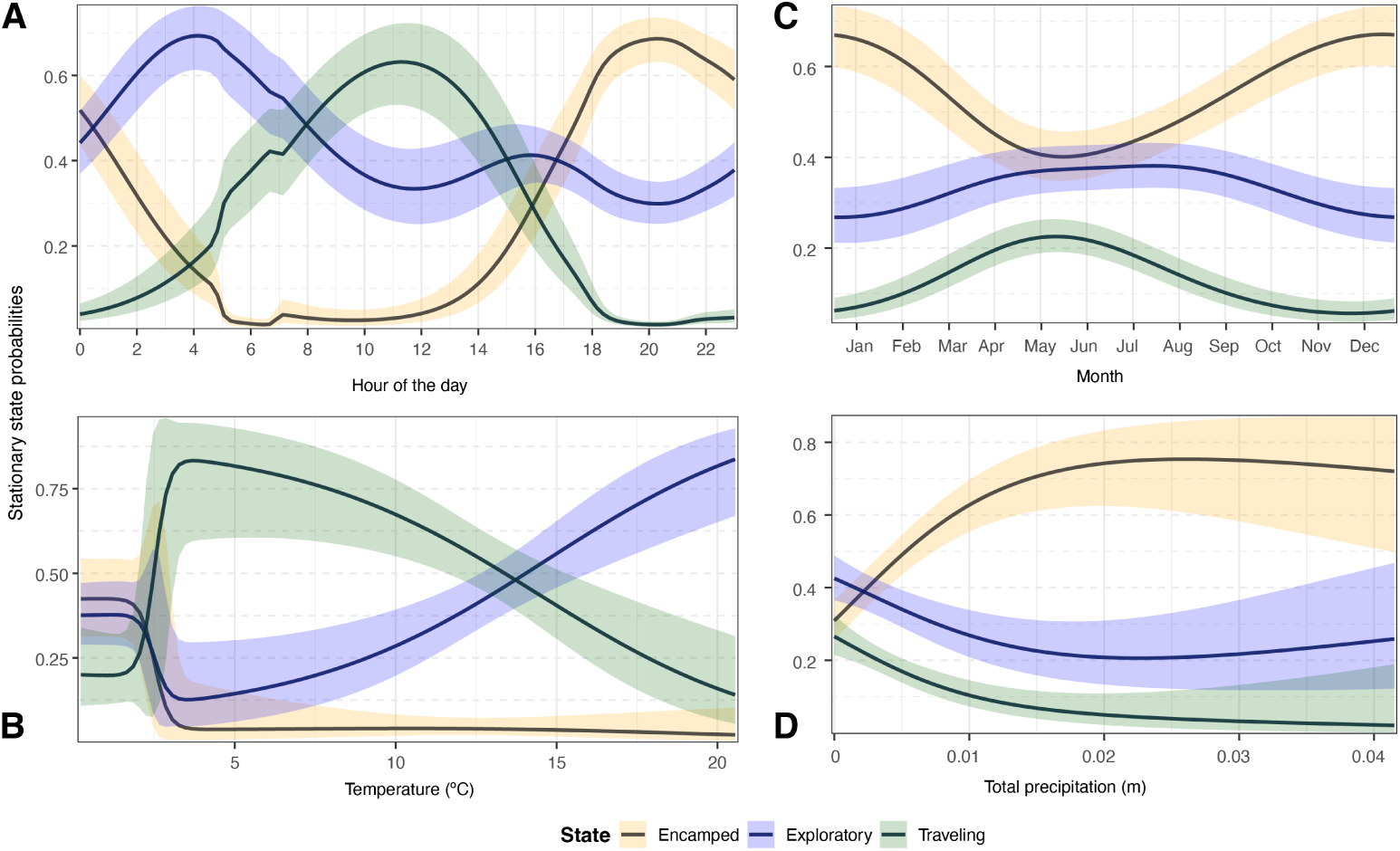
Stationary probabilities derived from a three-state Hidden Markov Model. The curved lines depict the estimated probability of the bears occupying each behavioral state as a function of day (A), temperature (B), day of the year (C), and precipitation (D) along with the 95% confidence intervals in shades. The inclusion of temperature and seasonal effects reflects the structure of the top-ranked model (ΔAIC-selected).

On the other hand, precipitation acted as a primary driver of seasonal behavioral transitions (Fig. 5C). The encamped state reached its maximum probability during the transition from late to early–year (October–January), suggesting a period of increased stationarity. Conversely, both foraging and large displacement probabilities peaked during sporadic rain and wet seasons (May–July), coinciding with a decrease in encamped behavior probabilities at 40%. The encamped state probabilities had a marked increase at higher rainfall levels as foraging and travelling decreased (Fig. 5D).

#### 3.2.2 Diel phenotypes

Hidden behind the traditional labels of diurnal or nocturnal, Andean bears exhibited a distinctly cathemeral activity cycle, as classified under the Traditional hypothesis, at the individual and population levels across behavioral states, sexes and regions (Fig. 6; Additional file 2, Table S9; Additional file 3, Fig. S6–S8).

**Fig. 6.**
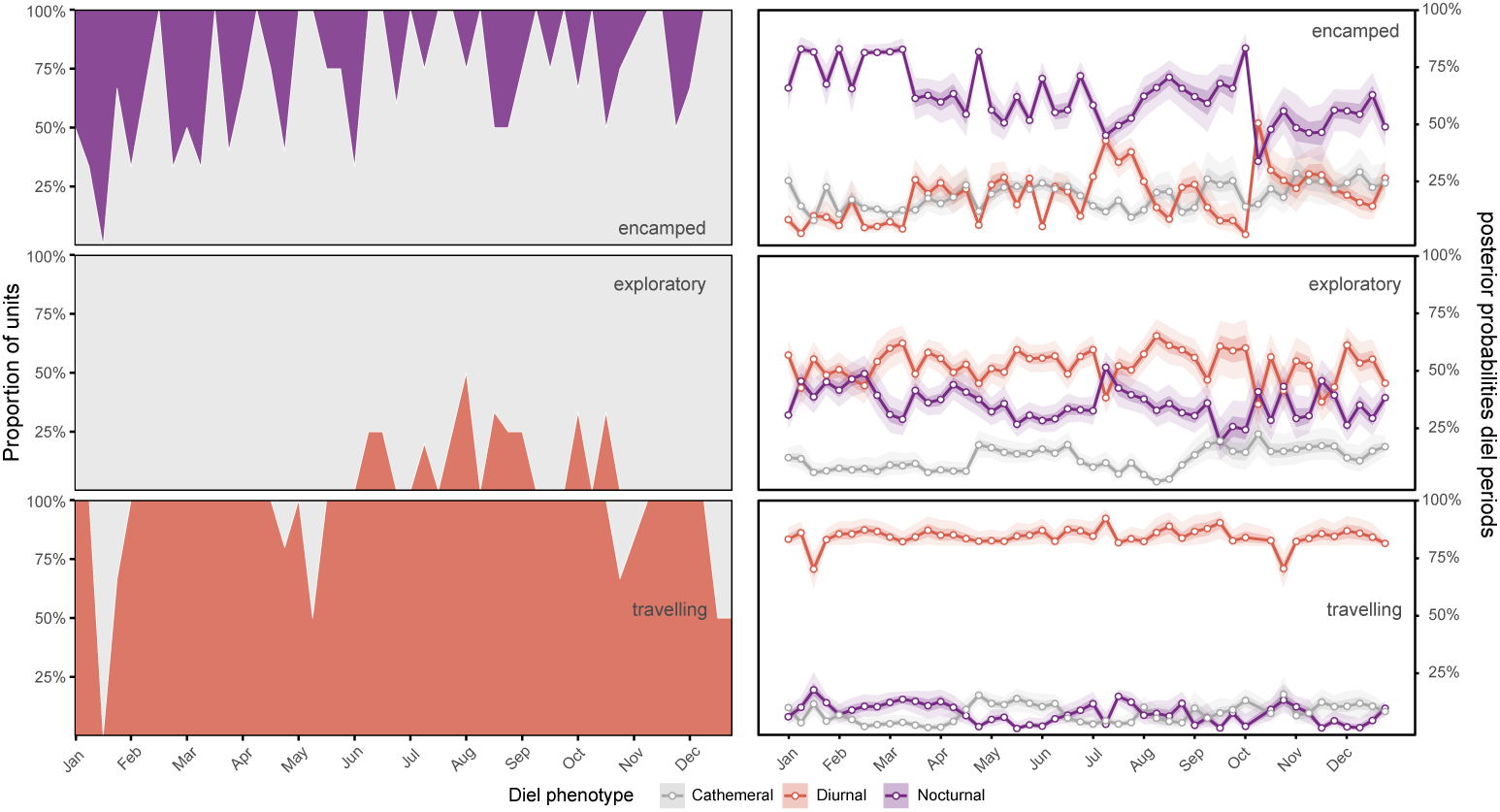
Diel phenotype weekly classifications under the Traditional hypothesis and their posterior probabilities across seasons and behavioral states. Left panels show the proportion of weekly units at the population-level classified as Nocturnal (purple), Cathemeral (gray), or Diurnal (orange). Right panels show posterior median probabilities of female activity in each diel period: nighttime (purple), twilight (gray) and daytime (orange), with dark to light shaded 95% and 50% credible intervals derived from Bayesian Markov Chain Monte Carlo sampling on the Diel.Niche activity distributions. Male patterns exhibit similar seasonal variation and equivalent posterior prob-abilities, see Additional file 3, Figs. S6–S9.

Males and females were uniformly cathemeral, although Matias returned two Diurnal classifications (Additional file 2, Table S9). When averaged across individuals, both Cayambe–Coca and Llanganates showed exclusive cathemeral classification, with no region showing diurnal preference (Additional file 3, Fig. S7).

Posterior median probabilities validated the phenotype classifications (Fig. 6; Additional file 3, Fig. S9–S10). Across behavioral states, sexes and regions, daytime and nighttime activity were both substantial while twilight activity remained the smallest component (posterior mean: *p*_tw_ ≈ 0.10 − 0.20; Fig. 6; Additional file 3, Figs. S8). Because neither the daytime nor the nighttime probability approached the 0.80 threshold above which a single-period phenotype is defined, the classifier returned a cathemeral phenotype in nearly every month. The travelling state, however, was the only exception: daytime activity rose to roughly 0.8–0.9, yielding predominantly Diurnal classifications (Fig. 6; Additional file 3, Figs. S8).

In the encamped state, nighttime activity was the largest component for most of the year (*p*_n_ ≈ 0.50 − 0.70), and in the individual months when it exceeded 0.80, the most-supported phenotype shifted to Nocturnal. Notably, this was more frequent and pronounced in females and in Llanganates (Additional file 3, Figs. S6–S7). Encamped activity was nighttime-weighted early in the year and shifted toward daytime later, but with no significant switches between diurnal and nocturnal phenotypes.

In the exploratory state, daytime and nighttime probabilities were more balanced (roughly 0.3–0.5), keeping the classification cathemeral apart from two sporadic Diurnal events in males when daytime activity was elevated (Additional file 3, Fig. S7).

Averaged across individuals, both Cayambe-Coca and Llanganates converged on daytime activity near 0.50 and nighttime near 0.40 with only minor twilight use, consistent with exclusive cathemeral classification and no diurnal preference at the regional level (Additional file 3, Fig. S9).

Detailed General and Selection hypotheses classifications are provided in Additional file 1. Briefly, similar patterns to the Traditional hypothesis were detected in the former; General Cathemeral and Diurnal-Nocturnal classifications were predominant at all levels and combinations, except for the travelling state and its combinations which were classified as Diurnal at proportions greater than 70%. Matias was the only bear to travel long distances at a 1:1 ratio of General Cathemeral and Diurnal phenotypes. The Selection hypothesis revealed a more complex pattern, with a greater diversity of classifications. However, the most common at different combinations was Day selection, followed by Day-Night and Equal selections (Additional file 1; Additional file 2, Table S9).

These diel phenotype classifications were independently corroborated by camera trap detections of untagged Andean bears from other localities in Ecuador, Colombia, and Peru [80]. These bears displayed behaviors traditionally seen during the day, including foraging, tree marking, and one observation of a male potentially chasing a female (Additional file 1). This suggests that the cathemeral pattern documented here could be a species-level trait rather than an artifact of individual behavior or collar scheduling, with rare strictly diurnal or nocturnal deviations.

### 3.3 Movement metrics

CTSD estimation was attempted at multiple temporal scales with feasibility that varied by individual and step size. Whole-tracking-period and diel estimates were restricted to GPS-tagged individuals, while weekly estimates were obtained for both tag types. At the whole-tracking-period scale, DeeAnn, Paya, and all VHF-tagged bears in Cotacachi-Cayapas (except for Fernando) were best described by an Ornstein-Uhlenbeck (OU) process (Additional file 2, Table S2), a CTMM inconsistent with speed estimation [73]. At the weekly scale, 328 out of 820 windows could not be estimated because sampling intervals exceeded the velocity autocorrelation timescale, with the highest failure rates in Matias (64), Chalpar (52) and Porraca (35), and additional 54 windows were excluded for having diffusion DOF *<* 5.

For diel-period estimates, average speeds were measurable for all bears during nighttime, though DeeAnn and Matias returned low ESS equal to 12 and 13, suggesting high measurement uncertainty (Additional file 2, Table S10). During diurnal periods DeeAnn and Paya’s average speeds remained inestimable for the same reason mentioned above, whereas Lorena exhibited the lowest *ESS* = 5 and wide CIs.

#### 3.3.1 Whole-tracking-period average speeds

Average speeds across the tracking duration revealed a population average of 6.04 km/day (5.08–7.13). While males moved slightly faster than females (6.37 vs. 5.64 km/day), this difference was not statistically significant (*p* = 0.08). Movement was more strongly influenced by geography; bears in Cayambe-Coca traveled significantly faster than those in Llanganates, averaging 6.74 km/day compared to 5.1 km/day (*p* = 0.003; Additional file 2, Table S11).

When isolating diurnal movement, activity levels varied significantly by age (Additional file 2, Table S11), adults were found to be more mobile than subadults (8.16 vs. 7.36 km/day; *p <* 0.001). Conversely, while nocturnal movement was generally lower, it was driven by both sex and region. Males were more active at night than females (1.63 vs. 1.19 km/night), and Cayambe-Coca bears nearly doubled the nocturnal speed of those in Llanganates (1.73 vs. 0.91 km/night; *p <* 0.00001).

Individually, Lorena and Matias exhibited the highest mobility during the day (11.19 km/day and 8.97 km/day, respectively), while the remainder of the cohort averaged approximately 7 km/day. However, Lorena’s CIs were wide (6 –16 km/day) due to a small ESS (Additional file 2, Table S10). Nighttime average speeds declined notably, yet Matias maintained the highest nocturnal speed at 2.71 km/night.

#### 3.3.2 Weekly decoupled speeds and diffusion rates

The speed and diffusion GAMs used all individuals except Fernando due to his low number of fixes. The speed model based on *n* = 426 weekly observations explained 81.2% of deviance, while the diffusion model based on *n* = 734 observations explained 45.3%, reflecting the greater robustness of diffusion estimation at coarse sampling intervals (Additional file 2, Table S12 and S13). Individual identity was a significant predictor in both models (*p <* 0.001), with diffusion showing greater among-individual variation than speed.

Speed and diffusion varied by region and season through different mechanisms. In the former, bears in Cotacachi-Cayapas moved substantially slower than those in Cayambe-Coca (*p <* 0.0001; Additional file 2, Table S12), with females moving around 11 times slower, yet speed was constrained in both sexes, hinting that dense vegetation may impose physical costs on movement. Seasonal variation in speed was nonlinear in Cotacachi-Cayapas and Llanganates (*p <* 0.001; Fig. 7), with no sex differences in the latter (*p* = 0.58; Additional file 2, Table S12). In contrast, diffusion showed complex seasonality only in Cayambe-Coca (*p <* 0.001) but no sex differences (*p* = 0.7; Fig. 7). A dissociation between speed and diffusion was observed in Llanganates (Fig. 7).

**Fig. 7.**
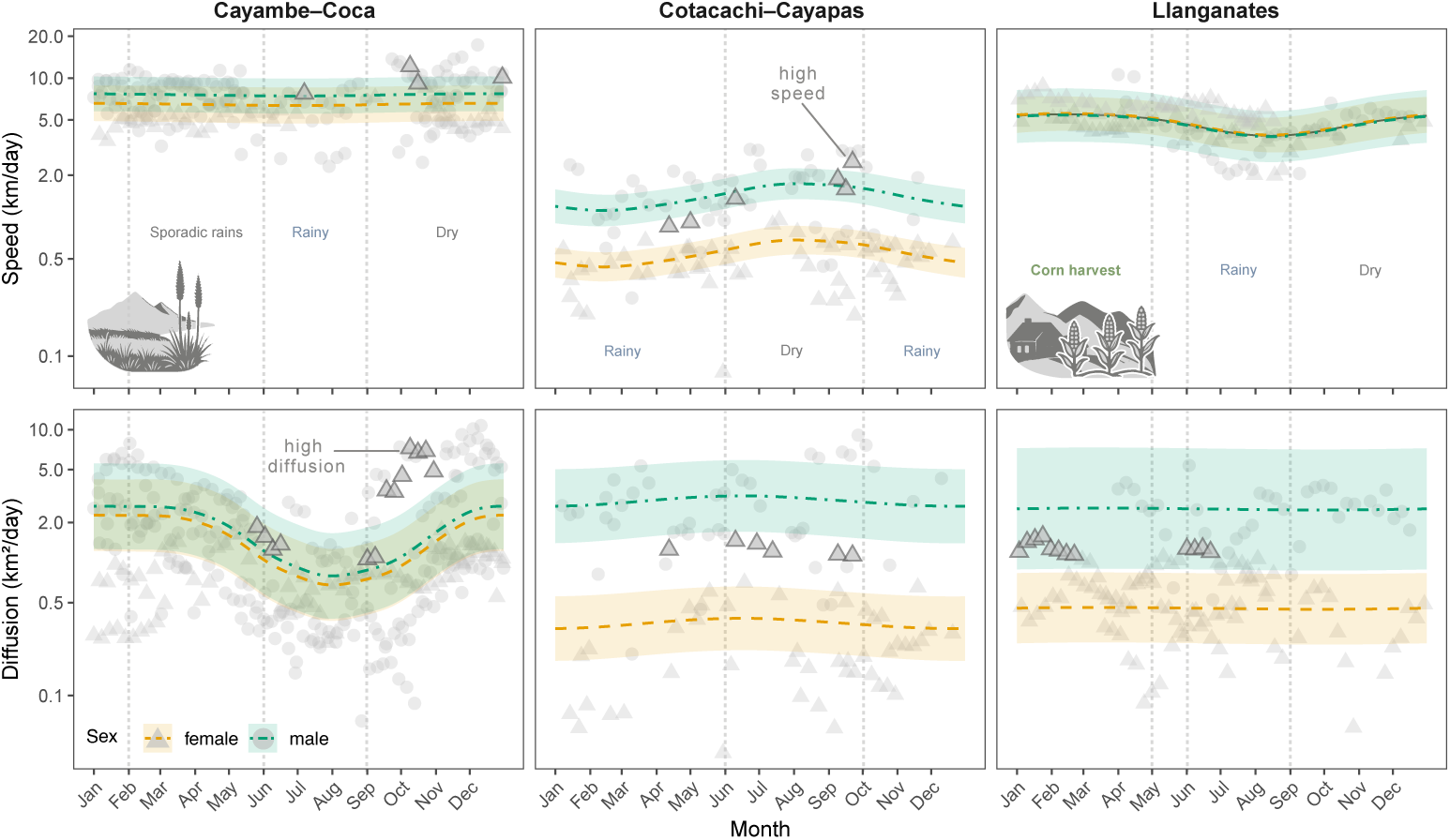
Seasonal variation in diffusion rates and average speeds by sex and region. Points show individual weekly estimates filtered with effective degrees of freedom *≥* 5. Colored lines and shaded ribbons show predicted means and 95% confidence intervals from GAMs fitted with a sex-region interaction. The y-axis is log_10_-scaled. Females with speed and diffusion higher than the regional average are labelled with dark gray triangles.

While speed in this NP was statistically indistinguishable from Cayambe-Coca (*p* = 0.18), spatial breadth was significantly reduced by approximately 66% (*p* = 0.014) and 73% in Cotacachi-Cayapas (*p* = 0.001). These findings suggest that bears in Llanganates move at a comparable pace but within more restricted space, consistent with the intensive use of localized resource patches noted in the home range analysis.

#### 3.3.3 Regional differences in intersex movement

Diffusion rates differed significantly between sexes in a region-dependent manner. Although the main effect of sex was non-significant in both models (*p >* 0.4; Additional file 2, Table S12 and S13), significant sex and region interactions revealed that vegetation context constrains female but not male ranging behavior. Female diffusion was strongly suppressed in Cotacachi-Cayapas (*p* = 0.001) and Llanganates (*p* = 0.013) relative to Cayambe-Coca where intersex ranging differences were insignificant. Male diffusion remained high or increased across regions (*p <* 0.05; Fig. 7).

In Cotacachi-Cayapas, males moved approximately 2 times faster than females (*p* = 0.003) and ranged around 7 times more broadly (*p* = 0.002), largely exceeding regional female metrics (Additional file 2, Table S12 and S13). In Llanganates, males showed marginally greater diffusion than females (*p* = 0.045) despite an absence of speed differences (*p* = 0.5; Fig. 7), reinforcing the distinction between the two metrics. Male bears in Llanganates covered more novel ground than females without moving proportionally faster, suggesting broader exploratory behavior rather than increased movement speed and therefore lower energy expenditure. This indicates that while locomotion speed is uniformly low in dense environments like Cotacachi-Cayapas for both sexes, area-use decisions remain more sex-strategy dependent.

#### 3.3.4 Instantaneous speeds

We observed both population and individual-level variation in instantaneous speeds, with diel cycles and sex emerging as primary variation sources. Based on 25,667 values, we detected spectacled bears moved faster during the day (median 0.3 km/h, range: 0.2–1.5) than at night (0.04 km/h, 0.01–0.7; MW: *p <* 0.0001; Fig. 8A; Additional file 3, Fig. S10). This diel contrast was further modulated by sex.

**Fig. 8.**
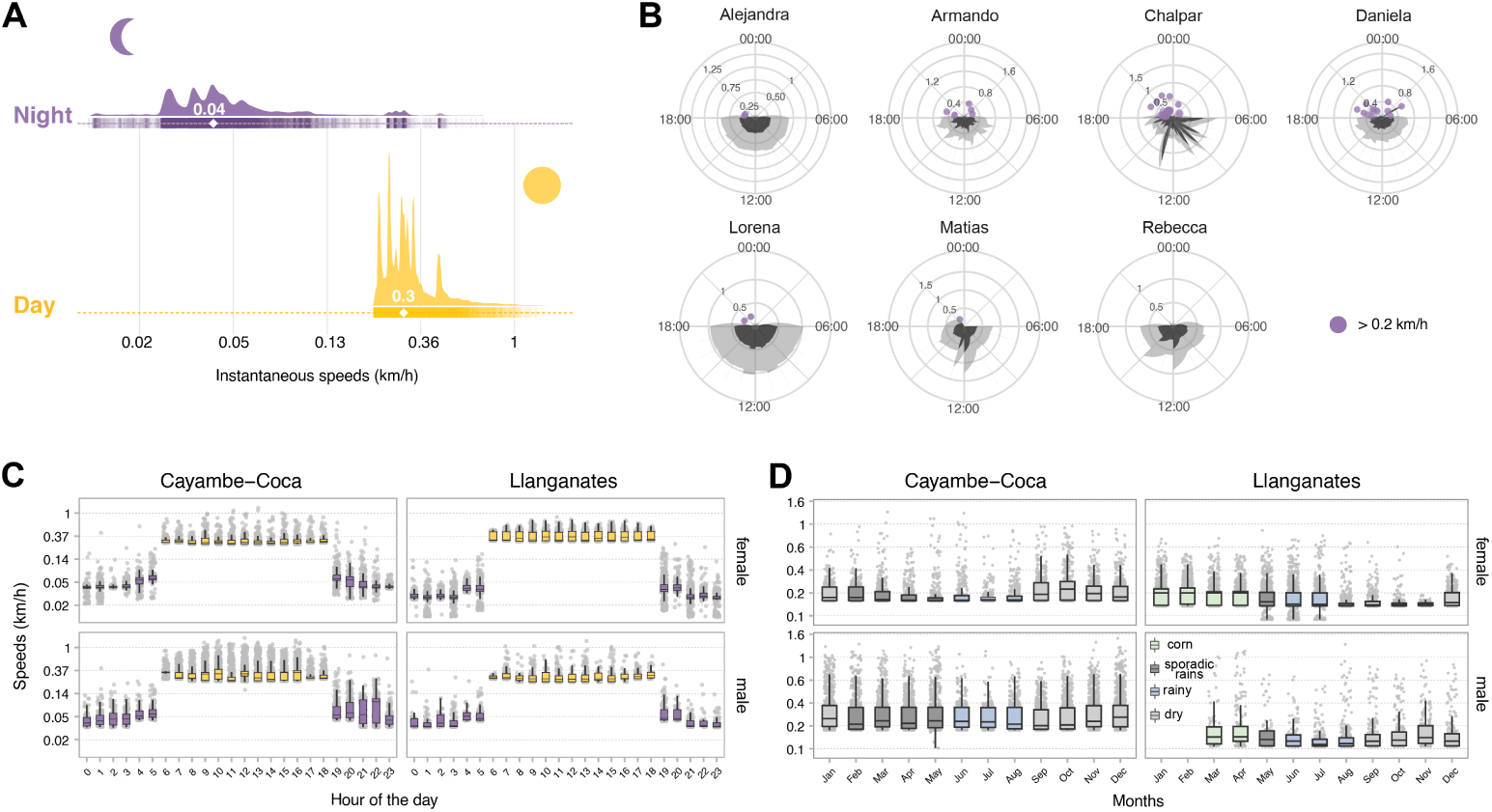
Continuous-time instantaneous speeds. (A) Raincloud plots show population-level diel-based speeds displayed on a log-transformed axis with ticks representing values on the natural scale. (B) Individual speeds (km/h) in 24-hour cycles. The 95% confidence intervals are shaded with dark and light grey indicating the lower and higher bounds of the estimate, respectively. Only nocturnal speeds *>* 0.2 are shown. (C) Boxplots of instantaneous speeds estimated from the diurnal and nocturnal subsets and for the (D) whole-tracking period displaying seasonal variation.

Males moved significantly faster than females (*p <* 0.0001) in a range of 0.03–1.5 km/h and displayed frequent speed bursts (Fig. 8B). Females traversed at speeds of 0.01–1.3 and moved slightly faster than males during daytime (*p <* 0.0001, *r* = 0.26; Additional file 3, Fig. S10). Individual variation was substantial within both sexes. During daytime hours, Chalpar reached up to 1.51 km/h, followed by Matias and Armando, both at 1.3 km/h. Both sexes accelerated beyond 0.14 km/h during nighttime, with fast-paced nocturnal bears peaking at 0.71 km/h for Chalpar, followed by Daniela (0.6 km/h), and Armando (0.4 km/h; Fig. 8B).

Habitat type significantly influenced movement speeds across diel cycles and seasons (Fig. 8C–D). Bears in Cayambe-Coca moved consistently faster than those in Llanganates (*p <* 0.001, Table 2). However, median speeds dropped significantly during the rainy season in both regions (KW: *p <* 0.0001). While males in Cayambe-Coca maintained seasonally constant speeds, females showed pronounced decelerations in the months of June–August (KW: *p <* 0.0001). Interestingly, the diel pattern shifted between parks: Llanganates bears were more active during the day, whereas Cayambe-Coca bears maintained higher relative speeds at night (*p <* 0.0001; Fig. 8C; Additional file 3, Fig. S11).

**Table 2.** Mann-Whitney U tests of instantaneous speeds. Median values (km/h) with observed minimum–maximum ranges in parentheses. All *p*-values represent pairwise region comparisons within each category: diel, region, sex, and season. Effect sizes are reported as rank biserial correlations (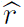), where positive values indicate higher speeds in Cayambe-Coca and negative show higher speeds in Llanganates.

| Category | Condition | Cayambe–Coca | Llanganates | $p$-value | $\hat{r}$ |
| --- | --- | --- | --- | --- | --- |
| Annual | Overall | 0.23 (0.14–1.51) | 0.18 (0.12–1.33) | $< 10^{-10}$ | 0.19 |
| Seasonal | Sporadic rains | 0.23 (0.19–1.33) | 0.22 (0.14–1.14) | $< 10^{-10}$ | 0.3 |
| | Rainy | 0.23 (0.14–1.27) | 0.17 (0.12–1.33) | $< 10^{-10}$ | 0.6 |
| | Dry | 0.23 (0.19–1.51) | 0.17 (0.14–0.92) | $< 10^{-10}$ | 0.7 |
| Sex | Female | 0.19 (0.18–1.30) | 0.18 (0.13–0.87) | 0.012 | 0.03 |
| | Male | 0.24 (0.14–1.51) | 0.16 (0.14–1.33) | $< 10^{-10}$ | 0.4 |
| Diel | Day | 0.28 (0.16–1.51) | 0.31 (0.22–1.32) | $< 10^{-10}$ | –0.3 |
| | Night | 0.05 (0.02–0.71) | 0.03 (0.01–0.48) | $< 10^{-10}$ | 0.7 |

#### 3.3.5 Movement intensities follow diel temperature cycles

The HGAM analysis explained 85.5% of deviance, revealing complex interactions driving movement speed and underlying the cathemeral activity pattern. The population-level diel smooth was highly significant (*p <* 0.0001), with predicted instantaneous speeds peaking between 12:00 and 16:00, declining through the evening, and a minimum between 00:00 and 05:00 (Additional file 3, Fig. S12). A secondary pre-dawn increase occurred between 05:00–07:00. Notably, speed did not approach zero at any hour, which is probably due to limitations with the CTMMs, which assume movement occurs at all times. While individual variation in these patterns was substantial (*p <* 0.0001), males exhibited significant seasonal deviations (*p <* 0.0001; Additional file 3, Fig. S13) that were minimal in females.

Temperature emerged as the primary environmental driver of diel activity (*p <* 0.0001;Additional file 2, Table S14; Additional file 3, Fig. S12). At thermal window of approximately 5°C–12°C, bears were substantially faster than average across most daytime and evening hours (08:00–22:00), particularly during the late afternoon (18:00–21:00; (Fig. 9A)). Conversely, at temperatures exceeding *>* 14°C, instantaneous speed was greatly suppressed across all hours, including during typically active periods. This heat avoidance behavior, was further evidenced by a modest positive partial effect detected during the pre-dawn window under warm conditions. Such flexibility allows individuals to exploit the coolest nightly hours when daytime heat surpasses their thermal tolerance.

**Fig. 9.**
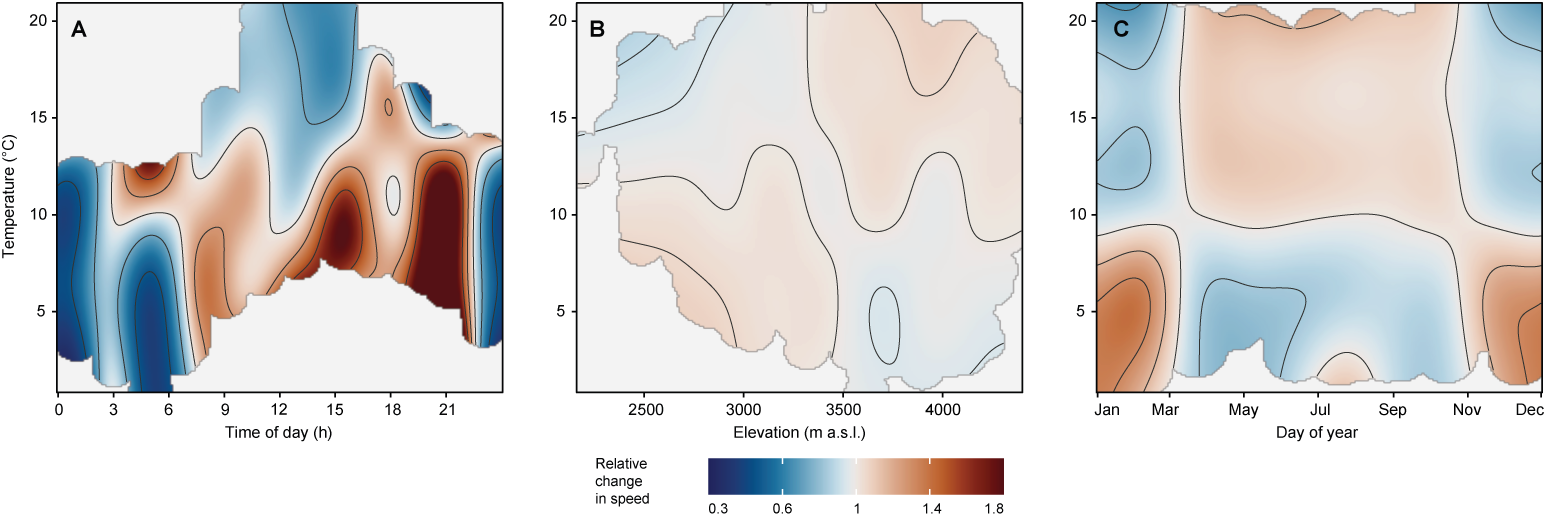
Interaction effects of temporal and environmental variables on instantaneous speeds. Heatmaps of the predicted partial effect on instantaneous speeds as a function of the temperature interaction with time of day (A), elevation (B), and day of year (C), expressed as the multiplicative change relative to the average partial effect of the interaction, where 1 indicates no deviation from the mean. Black contour lines in each panel are drawn at specific levels of multiplicative change. Light grey regions represent covariate combinations that do not occur in the empirical data.

Similarly, elevation and seasonality modified the relationship of temperature and speed. The interaction between elevation and temperature (*p <* 0.001) revealed that speeds increased above 3,500 m a.s.l. under warm conditions (Fig. 9B; Additional file 3, Fig. S12), indicating rapid and directed movements through the páramo. Likewise, cold temperatures at low elevations (*<* 2, 800 m a.s.l.) influenced rapid movements across agricultural and valley habitats during low-risk cool windows (Fig. 9B). Conversely, warm temperatures at low elevations produced the strongest movement suppression in the model.

The seasonal interaction with temperature was similarly well-supported (*p <* 0.0001), with cold conditions around November–March associated with enhanced movement, coinciding with the elevated speeds documented during the harvest and dry seasons in Llanganates, and warm mid-year temperatures producing broad movement suppression consistent with the rainy season diffusion contraction in Cayambe-Coca (Fig. 9C). The sampling interval correction term was significant at both the population and individual levels (*p <* 0.0001), confirming that GPS-derived speeds underestimate speeds at coarser sampling intervals [73] and that this bias varies across bears (Additional file 2, Table S14; Additional file 3, Fig. S13).

## 4 Discussion

Andean bears are increasingly venturing into the human domain in search of nourishment. While farming and cattle encroachment into natural habitats are well-known anthropogenic drivers of displacement, it is equally critical to understand the environmental factors shaping their movement behaviors, particularly under global warming scenarios. By compiling high resolution GPS data across multiple populations, this study achieved a fine-scale resolution that, until recently, was considered logistically unfeasible. This approach moves beyond broad range estimates to reveal the physiological and temporal constraints that dictate how Andean bears navigate a rapidly changing landscape.

### 4.1 Determinants of home range size

This first-of-its-kind population-scale analysis revealed sex as the main driver of home range and core area size, with males occupying territories, at least twice and in some cases up to fifteen times larger than those of certain females. For instance, Rebecca, who was rearing two cubs at a given time of tracking, exhibited restricted spatial use, likely reflecting strategies to minimize energy expenditure and risk during critical life stages [81], as documented in female grizzly bears [82, 83]. Nevertheless, females such as Porraca, Alejandra and Daniela exhibited areas comparable to those of males, suggesting exceptional spatial plasticity. At the individual-level, this demonstrates that females can forage across expansive landscapes as intensely as males, challenging traditional assumptions of sex-specific space use in the species.

Our home range estimates surpassed previous findings, including the reevaluated VHF-tagged bears [22], with the largest male reaching an average 369.9 km^2^, more than 150 km^2^ than the maximum home range reported in Colombia (238.86 km^2^ [23]). This discrepancy likely stems from area underestimation due to methodological differences (LoCoH vs AKDE, [28]), although a re-evaluation under the framework used here, could reveal underlying biological and ecological factors driving such variation.

Andean bears in this study traversed remote, rugged, and logistically challenging terrain. We found that weekly or seasonal displacements corresponded to frequent incursions into human-dominated landscapes, suggesting plastic and potentially adaptive movements across natural and modified environments (Additional file 1). These incursions suggest that habitat loss may be forcing individuals into human-modified environments due to widespread deforestation of native habitat [3, 84]. Our findings contrast with bear behavior in Colombia, where higher human density negatively affected bear presence [12, 85].

Early research hypothesized seasonal occupation of distinct habitats based on fruiting cycles in the Ecuadorian páramos [86], and Peruvian dry forests [87]. Our fine-scale analysis supports the existence of seasonal habitat use irrespective of sex, life stage, or region. Bears exhibited extended stationary behaviors in weekly or monthly periods, occupying reduced areas as small as 0.1–10 km^2^ (Fig. 3, Additional file 3, Fig. S3), likely driven by increased food availability [61]. In fact, young Chalpar remained for two months at the slopes of the Cayambe volcano (4,000 m.a.s.l.; Additional file 1), probably consuming *Hesperomeles obtusifolia*, a high-altitude (4,200 m.a.s.l) fruit that we have found to be part of the species diet in the area (Additional file 3, Fig. S14).

During corn harvest season in Llanganates, bears remained within reduced areas but moved away as the rainy season began (Fig. 3, Additional file 1), with larger displacements towards the páramos likely associated with the fruiting of mortiño. Similar range shifts occurred in Cayambe-Coca during both the rainy and dry seasons (Fig. 3), coinciding with the availability of achupalla in the páramos.

### 4.2 Spatial tolerance and overlap

Patterns of space sharing revealed how resource distribution and landscape fragmentation shape tolerance, yet with no evidence of close encounters between synchronously tagged individuals (Fig. 4; Additional file 1). As observed in Cotacachi-Cayapas, males and females showed near-complete spatial overlap while exploiting a mosaic of native vegetation and human-modified patches [27]. This included seasonal corn crops, perennial *Chusquea* bamboo stands, and epiphytic *Guzmania* bromeliads that are a source of water, sugar and protein year-round. Field observations confirmed that some females frequently occupied areas rich in achupallas, *Bromelia* spp., *Miconia*, and *Cavendishia*. Similarly, in Llanganates, overlaps between Armando, Alejandra, and the long-lived DeeAnn, were attributed to the seasonal availability of corn. The extensive area overlap appears to be driven by convergence on nutritional hotspots within fragmented landscapes across a steep altitudinal gradient (Fig. 4). Conversely, Lorena avoided the convergent territory and exploited other agricultural crops within reduced weekly home ranges (20.4 km^2^, Additional file 1), suggesting individual-level variation in foraging strategy.

These behaviors reinforce the hypothesis that Andean bear spatial ecology is shaped not only by intrinsic factors such as individual behavior, sex or evolutionary history (Additional file 1), but also by extrinsic drivers like resource stochasticity [61], and landscape structure. In fragmented Andean landscapes, where ecological richness coexists with anthropogenic pressure, spatial overlap may represent an adaptive strategy rather than evidence of habitat saturation.

### 4.3 Seasonal movement dynamics

While dynamic area estimates highlight how resources and fragmentation shape space use, movement metrics reveal the pace and scale at which bears navigate these landscapes, offering further insights into energy expenditure and landscape resistance (Additional file 1). Our analysis reveals that average speeds and diffusion rates capture different and complementary dimensions of Andean bear movement. The dissociation between these two metrics across study regions demonstrates that movement behavior in this species operates at multiple spatial and temporal scales, and corresponds precisely to the resource phenology of each protected area.

Speed and diffusion were suppressed in females in Cotacachi-Cayapas but diffusion sex differences were proportionally larger, with no evidence of seasonal fluctuations. Cotacachi-Cayapas is characterized by dense cloud forests at steep altitudinal gradients, with a mosaic of native vegetation and human-modified patches. The local landscape structure may, thus, impose substantial locomotion costs on patchy terrains [88], while simultaneously offering reliable, spatially concentrated food sources, reducing the need for extensive travel. This results in movement strategies characterized by slow, intensive, localized foraging rather than rapid and broad ranging. Male bears in the region reach diffusion rates comparable to males in Cayambe-Coca, which is consistent with typical space-use strategies driven by mate-searching as documented in brown bears [83].

The extensive home range and core area overlap between males and females documented in Cotacachi-Cayapas is thus consistent with movement patterns: females are moving slowly through resource-rich areas, while males range broadly over the same landscape at greater speed, with convergence occurring at predictable nutritional hotspots.

Males and females in Cayambe-Coca only display pronounced seasonal diffusion peaks during the dry seasons, coinciding with the availability of achupalla in the páramos [86]. This matches our home range analysis and supports hypotheses regarding bears tracking fruiting cycles [89]. The diffusion decrease during the rainy season suggests bears consolidate space use around reliable food sources and implies active habitat selection.

In Llanganates, speed seasonality was pronounced but diffusion seasonality was absent: speed increases during corn harvest without a corresponding increase in diffusion, which indicates intensive exploitation of localized agricultural patches (Fig. 7). The subsequent decline in speed during the dry season suggests a transition to lower-intensity foraging. This behavioral signature as fast and localized exploitation during the first months of the year, and slower and more exploratory thereafter has direct implications for managing agricultural exposure and human-wildlife conflict.

### 4.4 Sex biased dispersal and female plasticity

The spatial extents occupied by bears, coupled with the movement metrics presented here, elucidate a trade-off between energy expenditure and gene flow (Additional file 1), forming foundational hypotheses for future landscape genetics studies. Genetic evidence of female philopatry and restricted dispersal [11] aligns with the low diffusion rates (*<* 1 km^2^/day) and small core areas seen in females like Lorena and Paya. Conversely, males like Matias, Armando, and Pancho exhibited large home ranges and diffusion (2.7–4.1 km^2^/day), consistent with male-biased dispersal. Albeit Gilo had the largest areas and lowest male diffusion estimates across his tracking period (Additional file 2, Table S2).

Individual identity and plastic behaviors, thus, were shown to have a strong influence in diffusion rates, with some females exhibiting weekly movement metrics higher than males or the population average (Fig. 7). These deviations challenge a strictly sex-based dispersal model. Daniela, a female, demonstrated extreme displacement behavior, alternating between long-distance movements and localized residency. Similarly, Rebecca showed directionally persistent movement (Additional file 3, Fig. S5) with relatively high daytime diffusion rate (2 km^2^/day), and average speeds of 7 km/day, despite a compact core area (3.27 km^2^), suggesting episodic exploratory movement from a stable base. Showing that some bears are more spatially expansive than others regardless of sex, region, or season and are likely the primary drivers of long-distance gene flow and human-wildlife conflict.

This raises new hypotheses and challenges the traditional view of the sedentary Andean bear female: (1) female movement may be more plastic than previously assumed, especially in fragmented or resource-variable habitats suggesting that they might also play a critical role in maintaining population connectivity; and (2) speeds and diffusion rates may not solely reflect dispersal intent but also territorial negotiation or exploratory scouting, particularly in transitional life stages. Similar behaviors have been noted in giant pandas [90] and female brown bears [91], where this active role in mating is hypothesized as a means to avoid infanticide.

Future work integrating telemetry with genetic sampling could test whether high-diffusion females represent dispersing individuals or temporary range-shifters. Additionally, modeling landscape resistance in relation to female movement corridors may reveal whether philopatry is ecologically constrained or behaviorally reinforced.

### 4.5 Complex diel locomotion patterns

Movement metrics showed a marked increase in movement during daytime hours, with peak instantaneous speeds reaching 1.51 km/h and average speeds up to 8–12 km/day. Nevertheless, population estimates displayed notable nocturnal behavior of 1.54 km/night. Terrain steepness, however, did not appear to constrain movement speed: individuals such as Matias, Chalpar, and Lorena maintained high velocities despite consistently traversing steep slope landscapes even at nighttime hours given Matias’ displacements *>* 2 km/night (Fig. 8; Additional file 2, Table S10). These movement rates underscore the Andean bear’s capacity to sustain rapid travel across inclined terrain, a trait uncommon among ursids (Additional file 1).

Furthermore, diurnal and nocturnal movement patterns shifted between regions, bears in Llanganates were faster than those in Cayambe-Coca during the day. This regional variation may reflect differences in human activity patterns, and resource distribution. In fact, the HGAMs revealed that these differences are an emergent property of thermal switching and microclimatic conditions that influence the temporal diel niche of bears. The increased nocturnal speeds in Cayambe-Coca could also be a response to higher daytime human disturbance or a strategy to exploit nocturnal food resources.

### 4.6 Temperature-driven cathemerality

Andean bears reveal temporal flexibility in their activity schedules, underscoring that their movement ecology is shaped not only by spatial constraints but also thermal dynamics. This study offers the most detailed analysis to date of Andean bear movement patterns over a full 24-hour cycle, demonstrating that the species exhibit a cathemeral or diurnal-nocturnal phenotypes under General and Traditional hypotheses frameworks [72], while appearing mostly diurnal under Selection hypotheses sets.

This challenges previous assumptions that Andean bears are strictly diurnal [31] and reveals unexpected behavioral plasticity. While our dataset is limited to three Ecuadorian NPs and some camera trap records from international localities (see Additional file 1 [80]), the sporadic 24-hour activity patterns observed and supported by different analytical approaches, also align with observations from three reintroduced bears in Maquipucuna [34], located at least 100 km from the study sites.

The emergence of this pattern across disparate localities, thus, raises the question: is this a sustained behavior across the species’ geographical distribution or might this diel phenotype respond locally to unexplored environmental conditions? All other bear species have also been reported to be cathemeral (Additional file 1; [69, 92, 93]), hinting that this may be an ancestral character state widespread among the ursids, and which may have originated as a thermoregulatory adaptation that persisted across the bear radiation. By increasing movement during the night, Andean bears may inadvertently utilize a temporal window of opportunity to forage across anthropogenic landscapes when human surveillance is minimal. Several mammalian species have been documented to increase nocturnal habits as a strategy to circumvent human disturbance and persecution risk [94].

Both behavioral state prediction via HMMs and GAMs revealed that temperature is a key driver of diel behavioral switch and movement change. The appearance of activity distributed across the 24-hour cycle emerges not from a fixed internal preference but from temperature-driven behavioral responses. On cold days, bears maintained a broad diurnal-to-vespertine active window; on warm days, they suppress daytime locomotion and shift toward pre-dawn or evening activity.

Future research should explore how climate change-induced temperature shifts in Andean ecosystems may alter diel behaviors. Inferring how this diel phenotype will respond to drastic thermal fluctuations is an urgent priority. Furthermore, the relative influence of thermal constraints versus anthropogenic pressures remains to be assessed. Are bears actively avoiding humans at certain times of day, or is their nocturnal activity primarily driven by temperature? Understanding this balance is imperative for developing effective conservation strategies that mitigate human-wildlife conflict while also considering the impacts of climate change on bear behavior.

The Andean bear offers a particular study system for such investigation given the steep thermal gradients available across its altitudinal range, that we show to be relevant given its complex surface interaction with speed and temperature (Fig. 9).

### 4.7 Conservation strategies and recommendations

Compounding threats, including the increasingly frequent sightings of Andean bears near touristic infrastructure and highways [16], have contributed to its classification as Vulnerable on the IUCN Red List across its geographic range [95], and endangered in Ecuador since 2001 [96]. While past studies have noted human-bear interactions, this study highlights the urgent need for proactive and temporally dynamic conservation measures.

Our findings suggest that conservation strategies must reflect the extensive areas that both sexes occupy. The largest average male home range we documented (369.9 km^2^) raises concerns about whether current protected areas are sufficient to sustain viable populations. Similar challenges have been noted for other wide-ranging mammals like jaguars [97] and black bears [98], where protected areas alone are insufficient to encompass their ecological requirements.

Therefore, conservation strategies must account not only for broad home range requirements but also for diel and seasonal dynamics, and ecological context that influence how Andean bears use landscapes. The seasonal behavioral switches reported here—fast and localized during harvest, slower and more exploratory thereafter—have direct implications for human-wildlife conflict management, as they identify the first half of the year as the period of maximum agricultural exposure in regions like Llanganates and Cotacachi-Cayapas.

Given the dynamic and flexible nature of bear behaviors, such strategies must move beyond static mapping to incorporate the temporal dimensions of their ecology, addressing not only where risk is highest but also when and under what conditions bears are most active. However, to ensure these strategies are feasible, it is necessary to collaborate with local communities and organizations to verify if their implementation is actionable within the local socio-economic context. Furthermore, sustained engagement from government agencies remains essential to complement these efforts and ensure long–term protection of both bears and the ecosystems they help sustain. To this end, we propose: (1) creating ecological corridors accounting for multi-demographic movement that respond to behavioral plasticities and maintain movement continuity across fragmented forest patches; (2) prioritizing the designation of functional altitudinal linkages between highland and lowland ecosystems to mitigate human-wildlife conflict and safeguard crucial foraging grounds; (3) establishing dynamic buffer zones between agricultural areas and bear habitats that expand and are strictly enforced during peak seasonal activity to reduce direct encounters; (4) implementing community-based conservation programs that translate into actionable risk calendars for agro pastoral planning like grazing rotation or nocturnal livestock enclosure. Stakeholders can then mitigate conflict based on specific times of day or seasons when bears are most active. This ensures that conservation strategies address not only where bears move, but also when they are most likely to overlap with human activity.

### 4.8 Data sharing and collaboration

For decades, data sharing has been constrained by challenges in ensuring proper recognition for authors and collaborators. Field researchers often dedicate many years to collecting VHF, GPS, and camera trap data under physically demanding and resource-limited conditions, yet these datasets along with associated analytical code frequently remain unpublished. This “data lockdown” limits opportunities for independent reevaluation, slows regional comparisons and restrict the integration of emerging analytical tools.

Currently, collaborative platforms such as Movebank allow animal tracking data to be shared seamlessly while guaranteeing contributor attribution. These platforms have transformed research enabling cross-border analyses of movement ecology, habitat use, and conservation threats. For Andean bears, whose range spans multiple countries and diverse ecological zones, the potential for comparative synthesis is immense. We urge Andean bear researchers to accompany future publications with accessible datasets and reproducible code. Data sharing not only reduces research costs, but also accelerates, refines and broadens our collective understanding of this elusive species.

The long-term telemetry data presented here, spans over two decades and represents an investment of approximately USD 300,000, offering a foundational and invaluable resource for future research. By making these datasets accessible, we enable the reallocation of limited funding toward complementary conservation actions such as genetic sampling, landscape genetics, corridor modeling and community engagement. Our study demonstrates the value of sustained monitoring: detailed movement metrics, and diel activity patterns which reveal previously unknown behaviors. As technologies become more affordable and analytical methods more sophisticated, the potential to uncover the hidden lives of Andean bears grows rapidly. However, technological advancement alone is insufficient. Progress will depend on open data policies and cross-institutional collaboration within and across countries, ensuring that the knowledge gained from decades of fieldwork translates into tangible protection for the species and the ecosystems it sustains.

## 5 Conclusions

Individual bear home ranges among sexes are broader and more variable than traditionally assumed, with some females occupying areas as large as some males. Although there was a significant difference between sexes, the largest average home ranges are substantially larger than previously thought. Accordingly, conservation strategies should adopt a unified approach while enhancing the design and implementation of buffer zones which extend beyond existing protected areas to encompass the broader spatial needs of the species.

This study presents strong evidence supporting a cathemeral diel niche in a subpopulation of Ecuadorian Andean bears. While individual variation in movement patterns exists, the consistent influence of temperature as movement driver suggests that flexible diel behavior may be widespread across the species, supported by camera trap data from untagged specimens from Colombia, Ecuador and Peru. Future research should prioritize GPS-monitoring and camera trap integration across the species’ geographic range to further refine behavioral models by capturing twilight and nocturnal activity, a dimension historically underrepresented in ecological studies.

Finally, one of our central objectives is the democratization of ecological data. Openly sharing telemetry datasets not only honors the labor invested in long-term fieldwork but accelerates scientific progress and broadens conservation impact. This is urgent not only due to the rapid pace at which environmental change is transforming natural habitats into anthropogenic landscapes, but also as a mean to promote the training of future researchers and conservation practitioners. Regardless of the specific strategy, a fundamental pillar must be the strengthening of data collection efforts.

## Declarations

### Ethics approval and consent to participate

All capture and tracking efforts were conducted under permits issued by the Ministerio del Ambiente, Agua y Transición Ecológica del Ecuador (MAATE), including 004-11 IC-FAU-DPN/MA, 23-2014-IC-FAU-DPAP/MA, 23-2014-AD-IC-FAU-DPAP-MA, 003-2015-RIC-FAU-DPAP-MA, 003-2015-AD-RIC-FAU-DPAP-MA, 005-2017-RIC-FAU-DPAP-MA, 001-2018-IC-FAU-DPAP-MA, 001-2018-IC-AD-FAU-DPAP-MA, 004-2019-RIC-FAU-DPAPCH-MA, MAAE-ARSFC-2020-0642, MAAE-ARSFC-2021-1644, MAATE-ARSFC-2022-2583, and MAATE-ARSFC-2023-0145. To ensure minimal stress during immobilization, we followed standardized protocols for sedation, monitoring, and post-release evaluation, in accordance with best practices in wildlife research and the ethical guidelines of the American Society of Mammalogists’ Animal Care and Use Committee [99].

## Consent for publication

Not applicable.

## Availability of data and code

All code and files necessary to reproduce these results will be available upon publication at https://github.com/Xacfran/andean_bears. Only the fine-scale movement animations have been modified to exclude sensitive information like the longitude/latitude and dates to protect these vulnerable individuals. However, animations with sensitive information and the code to generate them can be shared with researchers or organizations upon request to the corresponding authors. Moreover, to ensure these findings are not locked behind a language barrier and hinder their practical application to conservation, we have released a Spanish version of this work also available on the Github repository.

## Declaration of Generative AI in scientific writing

During the preparation of this manuscript, FXC and DJ used Google Gemini 3.1 Deep Think models to assist with finding inconsistencies in the text, improving readability and suggesting additional references, which were read, reviewed, and validated before being added to cited literature. The authors reviewed and edited the content and take full responsibility for the content of this publication. Andean bear vectors in Figure 1 were generated from a sketch drawn by FXC and improved with Microsoft Designer and Adobe Illustrator Artificial Intelligences and further modified by FXC to keep biological accuracy.

## Author contributions

Conceptualization: FXC, AC. Data curation: FXC, AC. Formal Analysis: FXC. Funding Acquisition: AC, DJ, JB. Investigation: FXC, AC, DJ. Methodology: FXC, SM. Supervision: AC, SM. Validation: FXC, AC, SM. Visualization: FXC. Writing original draft: FXC, AC, DJ. Writing review & editing: all authors.

## Abbreviations

AKDE: Autocorrelated Kernel Density Estimate
CI: Confidence Interval
CTMM: Continuous-Time Movement Model
CTSD: Continuous-Time Speed and Distance
DOF: Degrees of Freedom
ESS: Effective Sample Size
GAM: Generalized Additive Model
GPS: Global Positioning System
HGAM: Hierarchical Generalized Additive Model
HMM: Hidden Markov Model
IID: Independent and Identically Distributed
KDE: Kernel Density Estimate
KW: Kruskal-Wallis Test
LoCoH: Local Convex Hull
MCP: Minimum Convex Polygon
MW: Mann-Whitney U Test
VHF: Very High Frequency

## Acknowledgements

We thank the team of guides and trackers whose tireless efforts made this work possible, including numerous park rangers and technicians. In the Cotacachi-Cayapas National Park, we were assisted by Milton Arcos, Alberto Tabango, and Samuel Ayala. In the Cayambe Coca National Park, we thank Melchor Ascanta, Rodrigo Ascanta, Óscar Ascanta, and their canine units— without whom the capture of large mammals in Ecuador would not be a reality. In Llanganates National Park, we thank the Martínez Pérez family, especially Edgar Martínez and Juan Fernando Duran.

We are grateful to all those who assisted directly or indirectly during the field expeditions: ángel Yánez and Leonardo Arias Cardenas. We especially thank Dolores and Gilberto Insuasti, Mariana Santos and Castellanos Peñafiel family for their unconditional support and encouragement through more than two decades of work. We are also grateful to Daniel Rodríguez, Danilo Vásquez, Heinz Plenge, and Francisco Sánchez Karste for kindly providing camera trap records as additional evidence of crepuscular and nocturnal activity in Andean bears.

This work was significantly improved by the teachings, insights, and recommendations of tutors and colleagues at the 2023 AniMove workshop. Likewise, Roland Kays and Natasja van Gestel guided early stages of the analysis, and Sean Sutor provided help with Figure 1, and suggested the inclusion of the HMMs to this work. We are indebted to you all.

Last but not least, we are profoundly grateful to Dave Goucher, Ivan Reyes, Fred and Donna Caswell, Michele Chartier, Jonathan Howarth, Ruthmery Pilco Huar-caya, Alan Roedell, Lorena Rodríguez, and two anonymous donors for their economic contributions toward the publication of this article.

## Additional files (available upon publication)

**Additional file 1.** Extended methods, results and discussion. Includes the description of weekly and monthly individual movements, and the description of crepuscular and nocturnal behaviors of untagged bears in Colombia, Ecuador and Peru captured by camera traps.

**Additional file 2.** Supplementary tables numbered in order of appearance in-text.

**Additional file 3.** Supplementary figures numbered in order of appearance in-text.

## References

[1] Velez-Liendo X, Jackson D, Ruiz-García M, Castellanos A. Andean Bear (Tremarctos ornatus). In: Bears of the World: Ecology, Conservation and Management. Cambridge University Press; 2020. p. 78–87.

[2] Garshelis DL. Family Ursidae (Bears). In: Wilson DE, Mittermeier RA, editors. Handbook of the Mammals of the World. vol. Vol. 1. Carnivores. Barcelona, Spain: Lynx Edicions; 2009. p. 448–497.

[3] García-Rangel S. Andean Bear *Tremarctos ornatus* Natural History and Conservation. Mammal Review. 2012;42(2):85–119. 10.1111/j.1365-2907.2011.00207.x.

[4] Vela-Vargas IM, Jorgenson JP, González-Maya JF, Koprowski JL. *Tremarctos ornatus* (Carnivora: Ursidae). Mammalian Species. 2021 Jul;53(1006):78–94. 10.1093/mspecies/seab008.

[5] Penteriani V, Melletti M, editors. Bears of the World: Ecology, Conservation and Management. 1st ed. Cambridge University Press; 2020.

[6] Ruiz-García M, Castellanos A, Arias-Vásquez JY, Shostell JM. Genetics of the Andean Bear (*Tremarctos ornatus*; Ursidae, Carnivora) in Ecuador: When the Andean Cordilleras Are Not an Obstacle. Mitochondrial DNA Part A: DNA Mapping, Sequencing, and Analysis. 2020;31(5):190–208. 10.1080/24701394.2020.1769088.

[7] Ruiz-García M, Castellanos A, Árias-Vásquez JY, Shostell JM. Effects of the Ecuadorian Andes on Genetic Structure of the Spectacled Bear with New Genetic Datasets. Scientific Reports. 2025 Sep;15(1):32735. 10.1038/s41598-025-17839-9.

[8] Ruiz-García M, Arias Vásquez JY, Restrepo H, Cáceres-Martínez CH, Shostell JM. The Genetic Structure of the Spectacled Bear (Tremarctos ornatus; Ursidae, Carnivora) in Colombia by Means of Mitochondrial and Microsatellite Markers. Journal of Mammalogy. 2020 Aug;101(4):1072–1090. 10.1093/jmammal/gyaa082.

[9] Ruiz-García M, Orozco-terWengel P, Castellanos A, Arias L. Microsatellite Analysis of the Spectacled Bear (*Tremarctos ornatus*) Across Its Range Distribution. Genes & genetic systems. 2005 Mar;80:57–69. 10.1266/ggs.80.57.

[10] Ruiz-García M. Molecular Population Genetic Analysis of the Spectacled Bear (*Tremarctos ornatus*) in the Northern Andean Area. Hereditas. 2003 Feb;138:81–93. 10.1034/j.1601-5223.2003.01578.x.

[11] Cueva DF, Zug R, Pozo MJ, Molina S, Cisneros R, Bustamante MR, et al. Evidence of Population Genetic Structure in Ecuadorian Andean Bears. Scientific Reports. 2024 Feb;14(1):2834. 10.1038/s41598-024-53003-5.

[12] Castrillón-Hoyos L, Rincón L, Troncoso-Saavedra J, Giraldo-Rojas M, Hernández-Rincón J, Velásquez-Vázquez A, et al. Occupancy and Habitat Use by the Andean Bear Are Negatively Affected by Human Presence and Forest Loss. Journal for Nature Conservation. 2023 Jun;73:126409. 10.1016/j.jnc.2023.126409.

[13] Morrell N, Appleton RD, Arcese P. Roads, Forest Cover, and Topography as Factors Affecting the Occurrence of Large Carnivores: The Case of the Andean Bear (Tremarctos ornatus). Global Ecology and Conservation. 2021 Apr;26:e01473. 10.1016/j.gecco.2021.e01473.

[14] Castellanos A. Maternal Behavior of a Female Andean Bear in the Paramo of Cayambe Coca National Park, Ecuador. International Bear News. 2015;24(1):32–33.

[15] Castellanos A, Jackson D. Does Rebecca, A Seasoned Andean Bear Mother, Show Seasonal Birthing Patterns? International Bear News. 2018;27(3):57–58.

[16] Castellanos A, Jackson D, Ascanta M. Are Reports of Cub Abandonment in Andean Bears a Result of Increasing Human Encroachment? International Bear News. 2019;28(1):14–15.

[17] Vasquez D, Castellanos A, Rivadeneira G. Social Behavior of the Andean Bear in Northern Ecuador. International Bear News. 2024;33(3):40–42.

[18] Castellanos A, Laguna A, Clifford S. Suggestions for Mitigating Cattle Depredation and Resulting Human-Bear Conflicts in Ecuador. International Bear News. 2011;20(3):16–18.

[19] Castellanos A, Medina MF, Beltran DF. Record of Canine Distemper Virus in an Andean Bear, Colombia. Boletín Técnico Serie Zoológica. 2023;18:5–8.

[20] Santodomingo A, Enríquez S, Thomas R, Muñoz-Leal S, Félix ML, Castellanos A, et al. A Novel Genotype of Babesia Microti-like Group in *Ixodes montoyanus* Ticks Parasitizing the Andean Bear (*Tremarctos ornatus*) in Ecuador. Experimental and Applied Acarology. 2025 Jan;94(2):30. 10.1007/s10493-024-00990-9.

[21] Castellanos A, Arias L, Ascanta M, Dadone L, Pukazhenthi B, Martínez E, et al. Mamíferos Grandes y Medianos En Ecuador: Recomendaciones Para Su Captura y Manejo. Mammalia aequatorialis. 2026;8(1):103–136. 10.59763/mam.aeq.v8i1.133.

[22] Castellanos A. Andean Bear Home Ranges in the Intag Region, Ecuador. Ursus. 2011 Apr;22(1):65–73. 10.2192/URSUS-D-10-00006.1.

[23] Rodriguez D, Reyes A, Tarquino-Carbonell AdP, Restrepo H, Reyes-Amaya N. Space Use by a Male Andean Bear (Tremarctos ornatus) Tracked with GPS Telemetry in the Macizo Chingaza, Cordillera Oriental of the Colombian Andes. Notas sobre Mamíferos Sudamericanos. 2021 Feb;3(1). 10.31687/saremNMS.21.2.4.

[24] Vela-Vargas IM, Arias-Bernal L, Koprowski J. Novel Insights into Andean Bear Home Range in the Chingaza Massif, Colombia. International Bear News. 2021;30(1):28–30.

[25] Paisley S, Garshelis DL. Activity Patterns and Time Budgets of Andean Bears (*Tremarctos Ornatus*) in the Apolobamba Range of Bolivia. Journal of Zoology. 2005;268:25–34. 10.1111/j.1469-7998.2005.00019.x.

[26] Pillco Huarcaya R, Whitworth A, Mamani N, Thomas M, Condori E. Through the Eyes of the Andean Bear: Camera Collar Insights into the Life of a Threatened South American Ursid. Ecology and Evolution. 2024;14(12):e70304. 10.1002/ece3.70304.

[27] Castellanos A. Andean Bear Core Area Overlap in the Intag Region, Ecuador. In: Molecular Population Genetics, Evolutionary Biology and Biological Conservation of the Neotropical Carnivores. Nova Science Publisers Inc; 2013. p. 531–543.

[28] Silva I, Fleming CH, Noonan MJ, Alston J, Folta C, Fagan WF, et al. Autocorrelation-Informed Home Range Estimation: A Review and Practical Guide. Methods in Ecology and Evolution. 2022;13(3):534–544. 10.1111/2041-210X.13786.

[29] Silva I, Fleming CH, Noonan MJ, Fagan WF, Calabrese JM. Too Few, Too Many, or Just Right? Optimizing Sample Sizes for Population-Level Inferences in Animal Tracking Projects. Ecology and Evolution. 2026;16(6):e73755. 10.1002/ece3.73755.

[30] Anand G, Fleming CH, Krishnan AG, Lamb CT, Medici EP, Prugh LR, et al. Estimating Population Range Distributions from Animal Tracking Data. bioRxiv. 2025 Sep;p. 2025.09.02.673746. 10.1101/2025.09.02.673746.

[31] Parra-Romero Á, Galindo-Tarazona R, González-Maya JF, Vela-Vargas IM. Not Eating Alone: Andean Bear Time Patterns and Potential Social Scavenging Behaviors. Therya. 2019;10(1):49–53. 10.12933/therya-19-625.

[32] Rodríguez D, Reyes-Amaya N, Reyes A, Restrepo H, Casas Y, Salgado O, et al. Desempeño de Un Collar GPS En El Seguimiento a Un Oso Andino (*Tremarctos ornatus*) En Los Andes Colombianos. Biodiversidad Neotropical. 2016 Mar;6:68–76. 10.18636/bioneotropical.v6i1.323.

[33] Peyton B. Spectacled Bear Conservation Action Plan. In: Servheen C, Herrero S, Peyton B, editors. Bears: Status Survey and Conservation Action Plan. IUCN, Gland, Switzerland, and Cambridge, UK; 1999. p. 157–193.

[34] Castellanos A, Altamirano BM, Tapi G. Ecología y comportamiento de osos andinos reintroducidos en la reserva biológica Maquipucuna, Ecuador: Implicaciones en Conservación. Politécnica. 2005;26(1):54–82.

[35] Ministerio del Ambiente. Plan de Manejo de la Reserva Ecológica Cotacachi Cayapas. Quito: Ministerio del Ambiente del Ecuador; 2007.

[36] Myers N, Mittermeier RA, Mittermeier CG, da Fonseca GAB, Kent J. Biodiversity Hotspots for Conservation Priorities. Nature. 2000 Feb;403(6772):853–858. 10.1038/35002501.

[37] Castellanos A. Guía para la rehabilitación, liberación y seguimiento de osos andinos. In: Castellanos A, Cevallos J, Laguna A, Achig L, Viteri P, Molina S, editors. Estrategia Nacional de Conservación del Oso Andino. Quito, Ecuador: Anyma; 2010. p. 1–34.

[38] Hijmans RJ, Cameron SE, Parra JL, Jones PG, Jarvis A. Very High Resolution Interpolated Climate Surfaces for Global Land Areas. International Journal of Climatology. 2005;25(15):1965–1978. 10.1002/joc.1276.

[39] Ministerio de Ambiente y Agua. Plan de Manejo del Parque Nacional Cayambe-Coca 2020-2030. Quito: Ministerio de Ambiente y Agua; 2020. Acuerdo Ministerial Nro. MAAE-2020-007. Available from: https://condesan.org/recursos/plan-manejo-del-parque-nacional-cayambe-coca-2020-2030/ [cited 2026 Jun 22].

[40] Sierra R. Propuesta preliminar de un sistema de clasificación de vegetación para el Ecuador continental. Quito, Ecuador: Proyecto INEFAN/GEF-BIRF y EcoCiencia; 1999.

[41] Albuja L, Almendáriz A, Barriga R, Montalvo L, Cáceres F, Román J. Fauna de Vertebrados Del Ecuador. Escuela Politécnica Nacional Instituto de Ciencias Biológicas; 2012.

[42] Varela LA, Ron SR. Geografía y clima del Ecuador. Quito: BIOWEB, Pontificia Universidad Católica del Ecuador; 2018. Available from: https://bioweb.bio/geografiaClima.html [cited 2026 Jun 20].

[43] Troya V, Cuesta F, Peralvo M. Food Habits of Andean Bears in the Oyacachi River Basin, Ecuador. Ursus. 2004 Apr;15(1):57–60. 10.2192/1537-6176(2004)015⟨0057:FHOABI⟩2.0.CO;2.

[44] Flores Velasco S, García JC. Environmental education as a tool for conservation of the spectacled bear (*Tremarctos ornatus*) in Oyacachi, Reserva Ecológica Cayambe-Coca, Ecuador. Lyonia. 2004;6(2).

[45] Ministerio del Ambiente. Plan de Manejo Parque Nacional Llanganates 2013. Quito: Ministerio del Ambiente del Ecuador; 2013. Available from: https://www.scribd.com/document/436815080/34-Plan-de-Manejo-Llanganates [cited 2026 Jun 22].

[46] Instituto Nacional de Meteorología e Hidrología. Anuario Meteorológico 2013. Quito, Ecuador: Instituto Nacional de Meteorología e Hidrología; 2017.

[47] Proyecto MapBiomas Ecuador. Colección 2.0 de la serie anual de mapas de cobertura y uso del suelo de Ecuador. MapBiomas Ecuador; 2023. Colección 2.0. Available from: https://ecuador.mapbiomas.org/ [cited 2025 Aug 21].

[48] Hollister J, Shah T, Nowosad J, Robitaille AL, Beck MW, Johnson M. Elevatr: Access Elevation Data from Various APIs. Comprehensive R Archive Network; 2023. R package version 0.99.0. Available from: https://github.com/jhollist/elevatr/.

[49] McDonald JE. Methods for Capturing Free-Ranging Black Bears, Ursus americanus, in Difficult Locations. The Canadian Field-Naturalist. 2003 Oct;117(4):621. 10.22621/cfn.v117i4.832.

[50] Allen RB. Research and Management Implications of the Pursuit of Black Bears with Trained Bear Dogs [Master of Science]. University of Montana. Montana, USA; 1985.

[51] Castellanos A, Arias L, Jackson D, Castellanos R. Hematological and Serum Biochemical Values of Andean Bears in Ecuador. Ursus. 2010 Jan;21(1):115–120. 10.2192/09GR002.1.

[52] Fleming CH, Drescher-Lehman J, Noonan MJ, Akre TSB, Brown DJ, Cochrane MM, et al. A Comprehensive Framework for Handling Location Error in Animal Tracking Data. bioRxiv. 2020 Jun;p. 2020.06.12.130195. 10.1101/2020.06.12.130195.

[53] Calabrese JM, Fleming CH, Gurarie E. ctmm: An r Package for Analyzing Animal Relocation Data as a Continuous-time Stochastic Process. Methods in Ecology and Evolution. 2016 Sep;7(9):1124–1132. 10.1111/2041-210X.12559.

[54] Castellanos FX, Jackson D, Mezzini S, Brito J, Castellanos A. Data from: Rethinking the Movement Ecology of Andean Bears: Temperature-Driven Cathemerality and Seasonal Space-Use Cycles. Movebank Data Repository; 2026. Available from: 10.5441/001/1.722.

[55] Fleming CH, Calabrese JM, Mueller T, Olson KA, Leimgruber P, Fagan WF. From Fine-Scale Foraging to Home Ranges: A Semivariance Approach to Identifying Movement Modes across Spatiotemporal Scales. The American Naturalist. 2014 May;183(5):E154–E167. 10.1086/675504.

[56] Alston JM, Fleming CH, Noonan MJ, Tucker MA, Silva I, Folta C, et al. Clarifying Space Use Concepts in Ecology: Range vs. Occurrence Distributions. Ecology. 2026;107(3):e70300. 10.1002/ecy.70300.

[57] Fleming CH, Sheldon D, Fagan WF, Leimgruber P, Mueller T, Nandintsetseg D, et al. Correcting for Missing and Irregular Data in Home-range Estimation. Ecological Applications. 2018 Jun;28(4):1003–1010. 10.1002/eap.1704.

[58] Fleming CH, Noonan MJ, Medici EP, Calabrese JM. Overcoming the Challenge of Small Effective Sample Sizes in Home-range Estimation. Methods in Ecology and Evolution. 2019 Oct;10(10):1679–1689. 10.1111/2041-210X.13270.

[59] Fleming CH, Deznabi I, Alavi S, Crofoot MC, Hirsch BT, Medici EP, et al. Population-Level Inference for Home-Range Areas. Methods in Ecology and Evolution. 2021 Jul;13:1027–1041. 10.1111/2041-210X.13815.

[60] Noonan MJ, Martinez-Garcia R, Davis GH, Crofoot MC, Kays R, Hirsch BT, et al. Estimating Encounter Location Distributions from Animal Tracking Data. Methods in Ecology and Evolution. 2021;12(7):1158–1173. 10.1111/2041-210X.13597.

[61] Mezzini S, Fleming CH, Medici EP, Noonan MJ. How Resource Abundance and Resource Stochasticity Affect Organisms’ Range Sizes. Movement Ecology. 2025 Mar;13(1):20. 10.1186/s40462-025-00546-5.

[62] Schwalb-Willmann J. Basemaps: Accessing Spatial Basemaps in R. Comprehensive R Archive Network; 2024. R package version 0.0.7. Available from: https://CRAN.R-project.org/package=basemaps.

[63] Castellanos FX, Jackson D, Mezzini S, Brito J, Castellanos A. Fine-scale spatiotemporal movements of Andean Bears in three Ecuadorian National Parks. Zenodo; 2026. Available from: 10.5281/zenodo.20821573.

[64] Dodge S, Bohrer G, Weinzierl R, Davidson SC, Kays R, Douglas D, et al. The Environmental-Data Automated Track Annotation (Env-DATA) System: Linking Animal Tracks with Environmental Data. Movement Ecology. 2013 Jul;1:3. 10.1186/2051-3933-1-3.

[65] Hersbach H, Bell B, Berrisford P, Biavati G, Horányi A, Muñoz Sabater J, et al. ERA5 hourly data on single levels from 1940 to present. Copernicus Climate Change Service (C3S) Climate Data Store (CDS); 2023. Available from: https://cds.climate.copernicus.eu/datasets/reanalysis-era5-single-levels [cited 2025 Sep 09].

[66] Muñoz Sabater J. ERA5-Land hourly data from 1950 to present. Copernicus Climate Change Service (C3S) Climate Data Store (CDS); 2019. Available from: https://cds.climate.copernicus.eu/datasets/reanalysis-era5-land [cited 2024 Apr 5].

[67] McClintock BT, Michelot T. momentuHMM: R Package for Generalized Hidden Markov Models of Animal Movement. Methods in Ecology and Evolution. 2018;9(6):1518–1530. 10.1111/2041-210X.12995.

[68] Revelle W. Psych: Procedures for Psychological, Psychometric, and Personality Research. Comprehensive R Archive Network; 2007. R package version 2.4.6. Available from: https://CRAN.R-project.org/package=psych.

[69] Kawamura K, Jimbo M, Adachi K, Shirane Y, Nakanishi M, Umemura Y, et al. Diel and Monthly Activity Pattern of Brown Bears and Sika Deer in the Shiretoko Peninsula, Hokkaido, Japan. The Journal of Veterinary Medical Science. 2022 Aug;84(8):1146–1156. 10.1292/jvms.21-0665.

[70] Lewis JS, Rachlow JL. Activity Patterns of Black Bears in Relation to Sex, Season, and Daily Movement Rates. Western North American Naturalist. 2011 Nov;71(3):388–395. 10.3398/064.071.0306.

[71] Donatelli A, Mastrantonio G, Ciucci P. Circadian Activity of Small Brown Bear Populations Living in Human-Dominated Landscapes. Scientific Reports. 2022 Sep;12(1):15804. 10.1038/s41598-022-20163-1.

[72] Gerber BD, Devarajan K, Farris ZJ, Fidino M. A Model-Based Hypothesis Framework to Define and Estimate the Diel Niche via the ‘Diel.Niche’ R Package. Journal of Animal Ecology. 2024;93(2):132–146. 10.1111/1365-2656.14035.

[73] Noonan MJ, Fleming CH, Akre TS, Drescher-Lehman J, Gurarie E, Harrison AL, et al. Scale-Insensitive Estimation of Speed and Distance Traveled from Animal Tracking Data. Movement Ecology. 2019 Nov;7(1):35. 10.1186/s40462-019-0177-1.

[74] DeNicola VL, Mezzini S, Cagnacci F, Fleming CH. Are Your Data Too Coarse for Speed Estimation? Diffusion Rates as an Alternative Measure of Animal Movement. bioRxiv. 2025 Jul;p. 2025.07.17.665364. 10.1101/2025.07.17.665364.

[75] Patil I. Visualizations with Statistical Details: The ‘ggstatsplot’ Approach. Journal of Open Source Software. 2021 May;6(61):3167. 10.21105/joss.03167.

[76] Wood SN. Generalized Additive Models: An Introduction with R. Second edition ed. Boca Raton, FL: CRC Press/Taylor & Francis Group; 2017.

[77] Pedersen EJ, Miller DL, Simpson GL, Ross N. Hierarchical Generalized Additive Models in Ecology: An Introduction with mgcv. PeerJ. 2019 May;7:e6876. 10.7717/peerj.6876.

[78] Wood SN, Goude Y, Shaw S. Generalized Additive Models for Large Data Sets. Journal of the Royal Statistical Society Series C: Applied Statistics. 2015 Jan;64(1):139–155. 10.1111/rssc.12068.

[79] Wood SN, Li Z, Shaddick G, Augustin NH. Generalized Additive Models for Gigadata: Modeling the U.K. Black Smoke Network Daily Data. Journal of the American Statistical Association. 2017 Jul;112(519):1199–1210. 10.1080/01621459.2016.1195744.

[80] Plenge H, Rodriguez D, Sánchez-Karste FJ, Vásquez D, Velazco R. Camera Trap Evidence of the Cathemeral Behavior of Andean Bears. Zenodo; 2026. Available from: 10.5281/zenodo.19964656.

[81] Séchaud R, Schalcher K, Almasi B, Bühler R, Safi K, Romano A, et al. Home Range Size and Habitat Quality Affect Breeding Success but Not Parental Investment in Barn Owl Males. Scientific Reports. 2022 Apr;12(1):6516. 10.1038/s41598-022-10324-7.

[82] Dahle, Swenson. Home Ranges in Adult Scandinavian Brown Bears (*Ursus arctos*): Effect of Mass, Sex, Reproductive Category, Population Density and Habitat Type. Journal of Zoology. 2003;260:329–335. 10.1017/S0952836903003753.

[83] Dahle B, Swenson JE. Seasonal Range Size in Relation to Reproductive Strategies in Brown Bears *Ursus arctos*. Journal of Animal Ecology. 2003;72(4):660–667. 10.1046/j.1365-2656.2003.00737.x.

[84] Kattan G, Hernández OL, Goldstein I, Rojas V, Murillo O, Gómez C, et al. Range Fragmentation in the Spectacled Bear *Tremarctos ornatus* in the Northern Andes. Oryx. 2004;38(2):155–163. 10.1017/S0030605304000298.

[85] González-Maya JF, Cáceres-Martínez CH, Acevedo Rincón A, Mauricio Vela-Vargas I. A True Highlander Hermit: Human Density and Distance to Natural Cover Negatively Affect Habitat Selection by Andean Bear. Journal for Nature Conservation. 2025 Mar;84:126833. 10.1016/j.jnc.2025.126833.

[86] Chávez AM, Díaz C, Amanzo JM. Seasonality of Andean Bear Scat Contents in Amazonas, Northeastern Peru. Ursus. 2021 Oct;2021(32e17):1–9. 10.2192/URSUS-D-20-00011.2.

[87] Suarez L. Seasonal Distribution and Food Habits of Spectacled Bears *Tremarctos ornatus* in the Highlands of Ecuador. Studies on Neotropical Fauna and Environment. 1988 Jan;23(3):133–136. 10.1080/01650528809360755.

[88] Halsey LG. Terrestrial Movement Energetics: Current Knowledge and Its Application to the Optimising Animal. Journal of Experimental Biology. 2016 May;219(10):1424–1431. 10.1242/jeb.133256.

[89] Peyton B. Ecology, Distribution, and Food Habits of Spectacled Bears, Tremarctos Ornatus, in Peru. Journal of Mammalogy. 1980 Dec;61(4):639–652. 10.2307/1380309.

[90] Connor T, Hull V, Liu J. Telemetry Research on Elusive Wildlife: A Synthesis of Studies on Giant Pandas. Integrative Zoology. 2016;11(4):295–307. 10.1111/1749-4877.12197.

[91] Penteriani V, Delgado MdM, Kojola I, Heikkinen S, Fedorca A, García-Sánchez P, et al. Mating from a Female Perspective: Do Brown Bear Females Play an Active Role in Mate Searching? Movement Ecology. 2025 Apr;13(1):24. 10.1186/s40462-025-00553-6.

[92] Sibarani MC, Ekanasty I, Surya RA. Using Bycatch Data to Model Sun Bear *Helarctos malayanus* Occupancy in Bukit Barisan Selatan National Park, Sumatra. Oryx. 2024 Jul;58(4):493–501. 10.1017/S0030605323001631.

[93] Ware JV, Rode KD, Robbins CT, Leise T, Weil CR, Jansen HT. The Clock Keeps Ticking: Circadian Rhythms of Free-Ranging Polar Bears. Journal of Biological Rhythms. 2020 Apr;35(2):180–194. 10.1177/0748730419900877.

[94] Gaynor KM, Hojnowski CE, Carter NH, Brashares JS. The Influence of Human Disturbance on Wildlife Nocturnality. Science. 2018 Jun;360(6394):1232–1235. 10.1126/science.aar7121.

[95] Velez-Liendo X, García-Rangel S. Tremarctos ornatus: The IUCN Red List of Threatened Species 2017: E.T22066A123792952. International Union for Conservation of Nature; 2016. e.T22066A123792952. Available from: https://www.iucnredlist.org/species/22066/123792952 [cited 2025 May 17].

[96] Cuesta F, Suárez L. Oso de anteojos. In: Tirira SD, editor. Libro rojo de los mamíferos del Ecuador. 1st ed. Serie Libros rojos del Ecuador. Quito, Ecuador: SIMBIOE; 2001. p. 68–70.

[97] de la Torre JA, Núñez JM, Medellín RA. Habitat Availability and Connectivity for Jaguars (*Panthera onca*) in the Southern Mayan Forest: Conservation Priorities for a Fragmented Landscape. Biological Conservation. 2017 Feb;206:270–282. 10.1016/j.biocon.2016.11.034.

[98] Zeller KA, Wattles DW, Conlee L, DeStefano S. Black Bears Alter Movements in Response to Anthropogenic Features with Time of Day and Season. Movement Ecology. 2019 Jul;7(1):19. 10.1186/s40462-019-0166-4.

[99] Sikes RS. 2016 Guidelines of the American Society of Mammalogists for the Use of Wild Mammals in Research and Education. Journal of Mammalogy. 2016;97(3):663–688. 10.1093/jmammal/gyw078.

